# A guidance into the fungal metabolomic abyss: Network analysis for revealing relationships between exogenous compounds and their outputs

**DOI:** 10.1101/2022.08.11.503656

**Authors:** Muralikrishnan Gopalakrishnan Meena, Matthew J. Lane, Joanna Tannous, Alyssa A. Carrell, Paul E. Abraham, Richard J. Giannone, Jean-Michel Ané, Nancy P. Keller, Jesse L. Labbé, David Kainer, Daniel A. Jacobson, Tomás A. Rush

**Affiliations:** National Center for Computational Sciences, Oak Ridge National Laboratory, Oak Ridge, TN 37831, USA; Biosciences Division, Oak Ridge National Laboratory, Oak Ridge, TN 37831, USA; Bredesen Center for Interdisciplinary Research and Graduate Education, University of Tennessee, Knoxville, TN 37916, USA; Department of Bacteriology, University of Wisconsin-Madison, Madison, WI 53706, USA; Department of Agronomy, University of Wisconsin-Madison, Madison, WI 53706, USA; Department of Medical Microbiology and Immunology, University of Wisconsin-Madison, Madison, WI 53706, USA; Now at Invaio Sciences, Cambridge, MA 02138, USA

**Author notes:** These authors contributed equally to this work (Muralikrishnan Gopalakrishnan Meena, Matthew J. Lane).

## Abstract

Fungal specialized metabolites include many bioactive compounds with potential applications as pharmaceuticals, agrochemical agents, and industrial chemicals. Exploring and discovering novel fungal metabolites is critical to combat antimicrobial resistance in various fields, including medicine and agriculture. Yet, identifying the conditions or treatments that will trigger the production of specialized metabolites in fungi can be cumbersome since most of these metabolites are not produced under standard culture conditions. Here, we introduce a data-driven algorithm comprising various network analysis routes to characterize the production of known and putative specialized metabolites and unknown analytes triggered by different exogenous compounds. We use bipartite networks to quantify the relationship between the metabolites and the treatments stimulating their production through two routes. The first, called the direct route, determines the production of known and putative specialized metabolites induced by a treatment. The second, called the auxiliary route, is specific for unknown analytes. We demonstrated the two routes by applying chitooligosaccharides and lipids at two different temperatures to the opportunistic human fungal pathogen *Aspergillus fumigatus*. We used various network centrality measures to rank the treatments based on their ability to trigger a broad range of specialized metabolites. The specialized metabolites were ranked based on their receptivity to various treatments. Altogether, our data-driven techniques can track the influence of any exogenous treatment or abiotic factor on the metabolomic output for targeted metabolite research. This approach can be applied to complement existing LC/MS analyses to overcome bottlenecks in drug discovery and development from fungi.

**Notice:** This manuscript has been authored by UT-Battelle, LLC, under contract DE-AC05-00OR22725 with the US Department of Energy (DOE). The US government retains and the publisher, by accepting the article for publication, acknowledges that the US government retains a nonexclusive, paid-up, irrevocable, worldwide license to publish or reproduce the published form of this manuscript, or allow others to do so, for US government purposes. DOE will provide public access to these results of federally sponsored research in accordance with the DOE Public Access Plan (http://energy.gov/downloads/doe-public-access-plan).

**Author summary:** Triggering silent biosynthetic gene clusters in fungi to produce specialized metabolites is a tedious process that requires assessing various environmental conditions, applications of epigenetic modulating agents, or co-cultures with other microbes. We provide a data-driven solution using network analysis, called “direct route”, to characterize the production of known and putative specialized metabolites triggered by various exogenous compounds. We also provide a “auxiliary route” to distinguish unique unknown analytes amongst the abundantly produced analytes in response to these treatments. The developed techniques can assist researchers to identify treatments or applications that could positively influence the production of a targeted metabolite or recognize unique unknown analytes that can be further fractionated, characterized, and screened for their biological activities and hence, discover new metabolites.

## Introduction

Fungi are among the most prolific producers of specialized metabolites classified by their chemical structure into four main classes: polyketides, non-ribosomal peptides, terpenes, and indole alkaloids [31, 30]. These specialized metabolites are multifaceted and impactful in our daily lives due to their positive roles as lifesaving drugs and agrochemicals. On the contrary, some of these specialized metabolites, commonly called mycotoxins, can have adverse effects on humans, animals, and crops, resulting in illnesses and economic losses [21, 43, 30]. In fungi, the genes involved in the biosynthesis of specialized metabolites are commonly arranged in so-called biosynthetic gene clusters (BGCs). Numerous specialized metabolite BGCs have been predicted and identified from genomes of several filamentous fungi [31, 30]. However, most predicted metabolites are not produced under standard cultivation and growth conditions, hindering their discovery [10]. In recent years, the chance to expand our knowledge and repertoire of specialized metabolites has significantly increased due to the enhanced understanding of fungal diversity and taxonomy, the widespread availability of published genomes [41], and the development of BGC prediction tools [11] such as antiSMASH [8, 7] and other computational tools. Biosynthetic genes predicted by these approaches can undergo genetic manipulations afterward to confirm their implication in the metabolic pathway and characterize novel specialized metabolites [45, 57]. However, this approach can sometimes be challenging, mainly if the BGC is missing a specific transcription factor to genetically target or if the gene cluster borders have been inaccurately predicted [54]. With the ongoing challenges of triggering those silent BGCs for specialized metabolite characterization, other approaches have been proposed.

The approaches adopted recently to trigger the expression of uncharacterized BGCs relied on identifying environmental cues, epigenetic chemical factors, axenic cultivation conditions, applications of exogenous compounds, or co-cultivation of the fungus with other microbes or hosts to induce the production of corresponding metabolites [37, 11, 35, 10, 49, 31, 30]. Network analysis [44] was also lately proposed as a complementary approach to accurately predict the factors that can elucidate fungal metabolites and narrow down the list of BGCs to target. In our recent paper, we demonstrated how the mathematical platform of graph theory [9] and network analysis [44] could target research to discover specialized metabolites within *Trichoderma* based on species-level taxonomic positioning and their predicted BCGs [54]. This study was later used to discover the antifungal agent, Ilicicolin H, in *Trichoderma reesei* [55]. There are various state-of-the-art techniques to help discover new specialized metabolites through network analysis, artificial intelligence, and data-driven approaches, like using molecular network analysis for web-based servers such as GNPS [62, 63], MetWork [4], and MetaboAnalyst [64, 48]. These web servers perform a variety of data-driven analyses on mass spectrometry data to discover new metabolites and characterize the structure of known and putative metabolites. Molecular networks are built using spectral matching (spectral network analysis [2]) to discover unknown compounds. However, to our knowledge, there are no tools developed to assess the direct effect of exogenous treatments on the production of fungal specialized metabolites. Thus, there is currently a void in the knowledge regarding the sources that trigger the production of such specialized metabolites.

Herein, our goal is to determine the feasibility of using network analysis to track the influence of applied exogenous compounds on the production of characterized and putative metabolites as well as unknown analytes. We introduce two methods based on network analysis to tackle these two objectives, called “direct route” and “auxiliary route”. An overview of the modeling framework is shown in Fig. 1 with suggestions of post-analysis applications. The direct route shows the influence of treatments on the production of known or putative specialized metabolites. In contrast, the auxiliary route distinguishes unique unknown analytes and pinpoint the treatments that foster their production. Both approaches reveal treatments that dominate by triggering a variety of specialized metabolites. Moreover, unique specialized metabolites are also identified by these methods. The capability of these methods was evaluated using the opportunistic human pathogen and soilborne saprotroph, *Aspergillus fumigatus*, as a model organism exposed to various chitooligosaccharides and lipid treatments.

**Figure 1:**
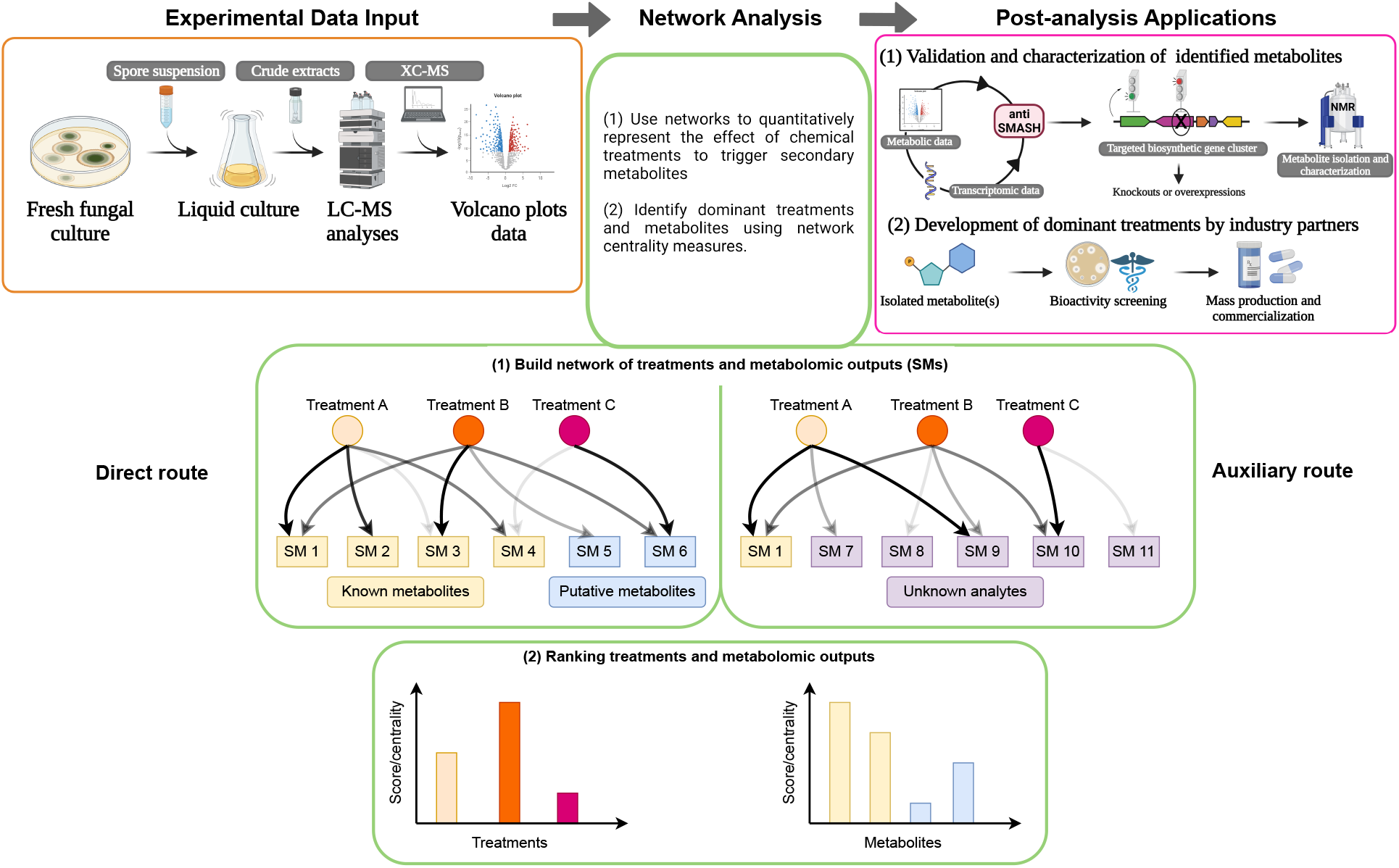
Framework of the direct and auxiliary routes using experimental inputs and implications for post-analysis applications. An overview of the network analysis approach reveals the effect of exogenous compounds on triggering the production of microbial specialized metabolites. **Experimental data input:** Experimental data from LC/MS analysis is used to reveal significant analytes. [54] provides a detailed roadmap on how to derive the experimental data. **Network analysis:** Network analysis processes the LC/MS data points to define graphs which quantitatively represent the effect of exogenous treatments to trigger specialized metabolites. The direct route is used to understand the influence of treatment on known or putative metabolites. In addition, after the data is curated to eliminate peak noise, the auxiliary route is used to identify strong signals of unknown analytes. The treatments and metabolomic outputs are ranked using network analysis measures. **Post-analysis applications:** After knowing the relationship between a treatment and metabolomic outputs, post-analysis applications can be applied to isolate and characterize the metabolite through genetic manipulations followed by bioactivity screening. These two post-analysis applications are provided as a guidance on possible applications of the current framework for discovering new metabolites and are not performed in the current study.

We emphasize that the particular objective of the current study is not to discover new metabolites whereas is to reveal the effect of exogenous treatments on triggering the production of specialized metabolites. Nonetheless, the inferences drawn on the dominant treatments and unique specialized metabolites using the current methodology can enable the discovery of new metabolites through the suggested post-analysis applications in Fig. 1 (top-right box). The developed methods can complement existing web-based network analysis tools [62, 63, 4, 64, 48] for metabolite discovery that do not consider the sources triggering metabolomic outputs and have great application potentials in various fields including drug discovery and development.

## Framework to provide data-driven network analysis

### Methodology: Building bipartite networks of treatments and metabolomic outputs

Bipartite networks are built to quantify the relationship between metabolites and the sources triggering their production, such as various exogenous biomolecules or compounds. A network (or graph) is a collection of nodes connected by lines named edges. The nodes represent the entities or elements of a system, and the edges represent the interaction or relationship amongst the features [9, 44]. For example, in cell metabolism, a metabolic network represents the biochemical reactions amongst substrates that result in products [26, 61, 44]. The nodes of the metabolic network represent the substrates, and the edges represent the metabolic reactions amongst the substrates. In the current analysis, we assessed the effect of exogenous compounds on the production of specialized microbial metabolites. This relationship between treatments and specialized metabolites can be represented by a network, as shown in Fig. 1. The nodes can be classified into two types: the treatments and the specialized metabolites, resulting in a bipartite network. The edges represent the magnitude of up- or down-regulation of specialized metabolites caused by the treatments compared to a controlled case (measured by the magnitude of log_2_ fold change using processed spectral data from targeted LC-MS analysis). The details of building the bipartite network are provided in the **Materials and methods** section.

The bipartite network provides an in-depth quantification and clear visual representation of a treatment’s ability to trigger the production of various specialized metabolites. We provide two routes to assess specialized metabolite production using the bipartite network formulation, as shown in Fig. 1. The first is the direct route to determine the biosynthesis of known and putative metabolites, whereas the second is the auxiliary route to assess the production of unknown analytes. In the direct route, the network nodes include treatments that elucidate known and putative metabolites by a microbe. Moreover, we use network centrality measurements to rank the treatments and the specialized metabolites. Those measurements are usually used to identify the most influential nodes in a network [44]. Herein, we use the centrality measurements of node strength and PageRank [12] to identify the most effective treatments and metabolites. The treatments are ranked based on their capability to trigger metabolite production, and the metabolites are classified based on their popularity in being activated by various treatments. We provide details of computing the network centrality measures in the **Materials and methods** section. In the auxiliary route, we built bipartite networks using novel analyte peaks extracted from post-processed spectral data. Furthermore, we analyze the edges and neighborhoods of nodes to distinguish unique analytes among the total pool. Methods for novel peak selection are provided in the **Materials and methods** section. Note that the only similarity between the two routes is on the definition of the graphs/networks - both involves bipartite networks defined by the amount of up- or down-regulation of the specialized metabolites by the exogenous treatments. The data curation and network centrality measures used to identify important specialized metabolites and treatments in each of the routes are different.

### System: Experimental data of *Aspergillus fumigatus* metabolomic outputs

Our modeling framework was used to reveal the effect of various chitooligosaccharides and lipid treatments on triggering the production of specialized metabolites in *Aspergillus fumigatus*. Various chitooligosaccharides and lipids were applied as exogenous treatments since they are common constituents found in most fungi [42, 53, 15]. Moreover, it has been previously shown that lipids influence fungal metabolomics [40]; however, the impacts of chitooligosaccharides have remained unknown. In contrast, chitooligosaccharides are reported to have antifungal activity [34], which might potentially influence the metabolomic profile in *Aspergillus* species [37]. We applied the treatments to *A. fumigatus* strain Af293 because it has a well-defined repertoire of known and putative specialized metabolites [52] that is temperature-dependent [32, 37, 5, 35, 24] and its whole genome is sequenced [46]. We also explore the influence of temperature on the production of specialized metabolites conducting the experiments at 25 and 37°C. *Aspergillus fumigatus* is generally examined at 25°C to explore the extent of its metabolomic capabilities or its lifestyle as a soilborne saprotroph that recycles environmental carbon and nitrogen [33]. However, the fungus is also an opportunistic human pathogen and is commonly examined at 37°C for its ability to cause aspergillosis, a lung disease found in immunocompromised patients [14]. The details of the experimental setups and data generation are provided in the **Materials and methods** section.

## Results

### Data-driven validation of direct and auxiliary routes with treated samples at 25°C

#### Analyte and metabolomic production induced by treatments

We use UpSet plots (Fig. 2a) and volcano plots (Figs. 2b-f) to curate the analytes and metabolites produced. The data curation for the UpSet plots (Fig. 2a and Fig. 5a for results of 37°C) and volcano plots (Figs. 2b-f and 5b-f for results of 37°C) were obtained using the experimental results involving XCMS [17] to provide a statistical assessment of signals with significant peaks between a treatment and solvent control through a pairwise comparison. A list of mass to charge (m/z) and retention times and statistical values were provided from those results. To validate the XCMS results, we used MAVEN [39, 13] for metabolomic analysis and visualization of the LC/MS data to confirm if the same m/z at the provided retention time did have a significant peak difference between treatment and solvent control.

**Figure 2:**
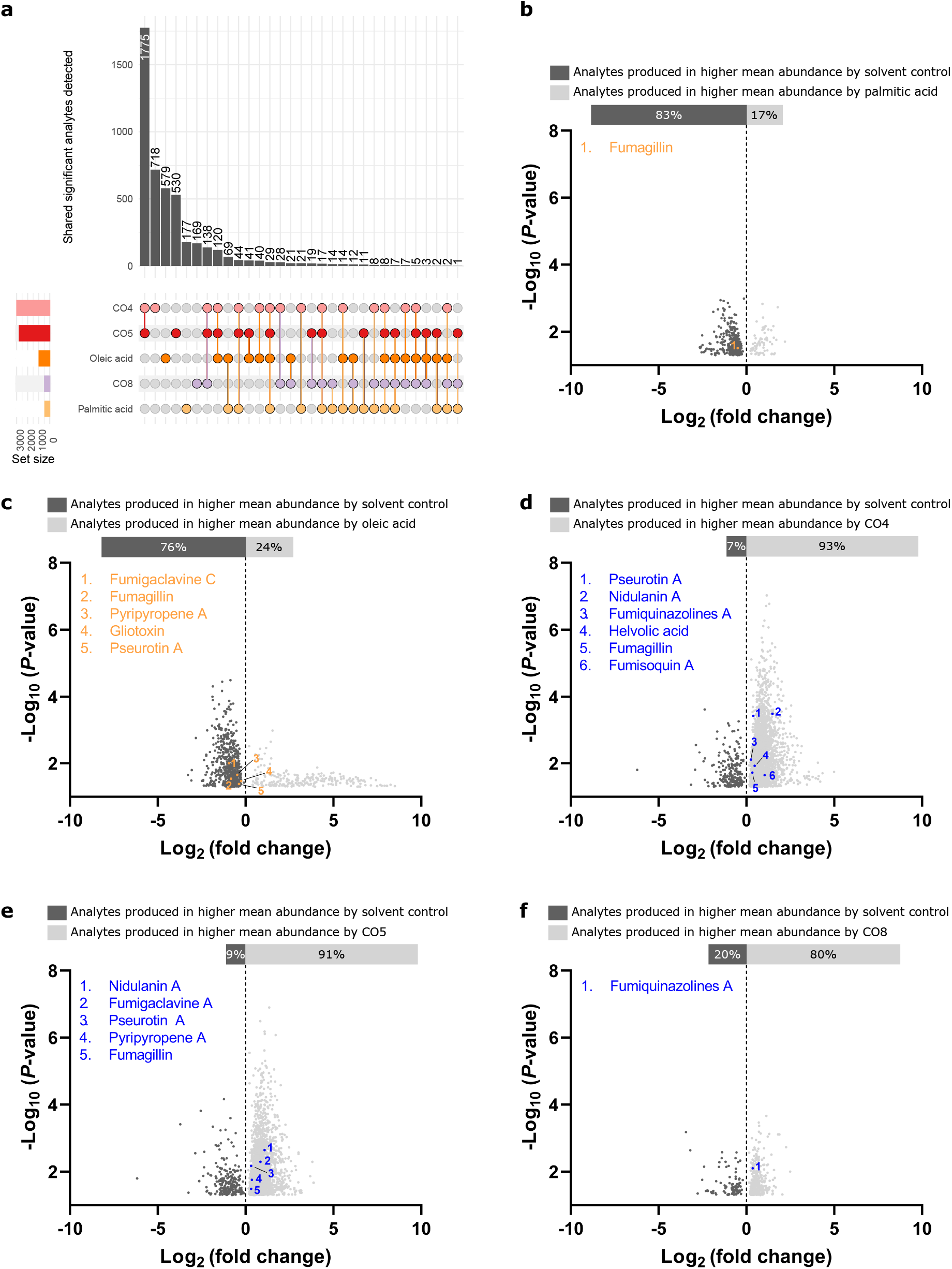
Metabolomic outputs of *Aspergillus fumigatus* at 25°C induced by lipid and chitooligosaccharide treatments. (a) UpSet plot denoting the number of significant analytes produced by individually applied treatments. Multiple treatments induced some analytes. (b-f) Volcano plots identifying the known and putative metabolites and unknown analytes triggered by (b) palmitic acid, (c) oleic acid, (d) CO4, (e) CO5, and (f) CO8 as compared to the solvent control.

At 25°C, LC/MS data revealed that all individually applied treatments significantly induced the production of analytes compared to the solvent control, as shown by the UpSet plot in Fig. 2a. No treatments were co-applied to the fungus.

In total, 4,629 significant analytes were detected. (Fig. 2a). Unique analytes produced by CO4 was 15.5%, by CO5 was 11.4%, by CO8 was 3.7%, by palmitic acid was 3.8%, and by oleic acid was 12.5%. These results indicate the length of COs matters when looking at how they influence specific metabolomic pathways. The short-chain COs has higher production of metabolites compared to long-chain COs. In addition, oleic acid has a more significant impact on analyte production compared to palmitic acid. Interestingly, 38.3% of individual analytes were induced by CO4 and CO5, suggesting a common co-regulation of metabolomic pathways. However, most analytes were uniquely caused by one treatment indicating that metabolomic pathways seem specific to treatment, thus requiring network analysis to determine those relationships.

To determine the regulation of analytes induced by treatment and identify potential known and putative specialized metabolites, we made volcano plots based on the log_2_ fold change and − log_10_ (p-values) between a treatment and the solvent control shown in Figs. 2 b-f. All treatments had a significant differential expression of analytes compared to the control. COs induce the production of analytes between 80% to 93% compared to the solvent control, including several known and putative metabolites (Figs. 2 d-f). On the contrary, the positive regulation of analyte production by lipids was reduced compared to the control, ranging between 17% to 24%. CO4 (Fig. 2d) and CO5 treatments (Fig. 2e) induced the production of five to six known or putative metabolites, whereas CO8 (Fig. 2f) influenced the production of a single known metabolite. These results follow the same trend shown in Fig. 2a, highlighting that short-chain COs have a more significant impact on triggering metabolite production than long-chain CO. Interestingly, oleic acid (Fig. 2c) induced five times more known or putative metabolites than palmitic acid (Fig. 2b), matching the data shown in Fig. 2a. We also compared transcriptomic expression from quantitative PCR analysis to corrected peak area which so relatively similar regulation of known secondary metabolites (Fig. 3). It is important to consider that specialized metabolomic production is potentially subject to post-transcriptional regulation when the transcriptomic and metabolomic data do not match. Discrepancies between metabolomic and transcriptomic profiles have been observed in other ascomycete fungi [56]. In summary, short-chain COs and oleic acid have the most significant impact on known or putative metabolite and unknown analyte production, yet they have different regulations at 25°C. The direct route using theoretical graph analysis is applied to understand further this regulation and how these metabolites are produced within a system.

**Figure 3:**
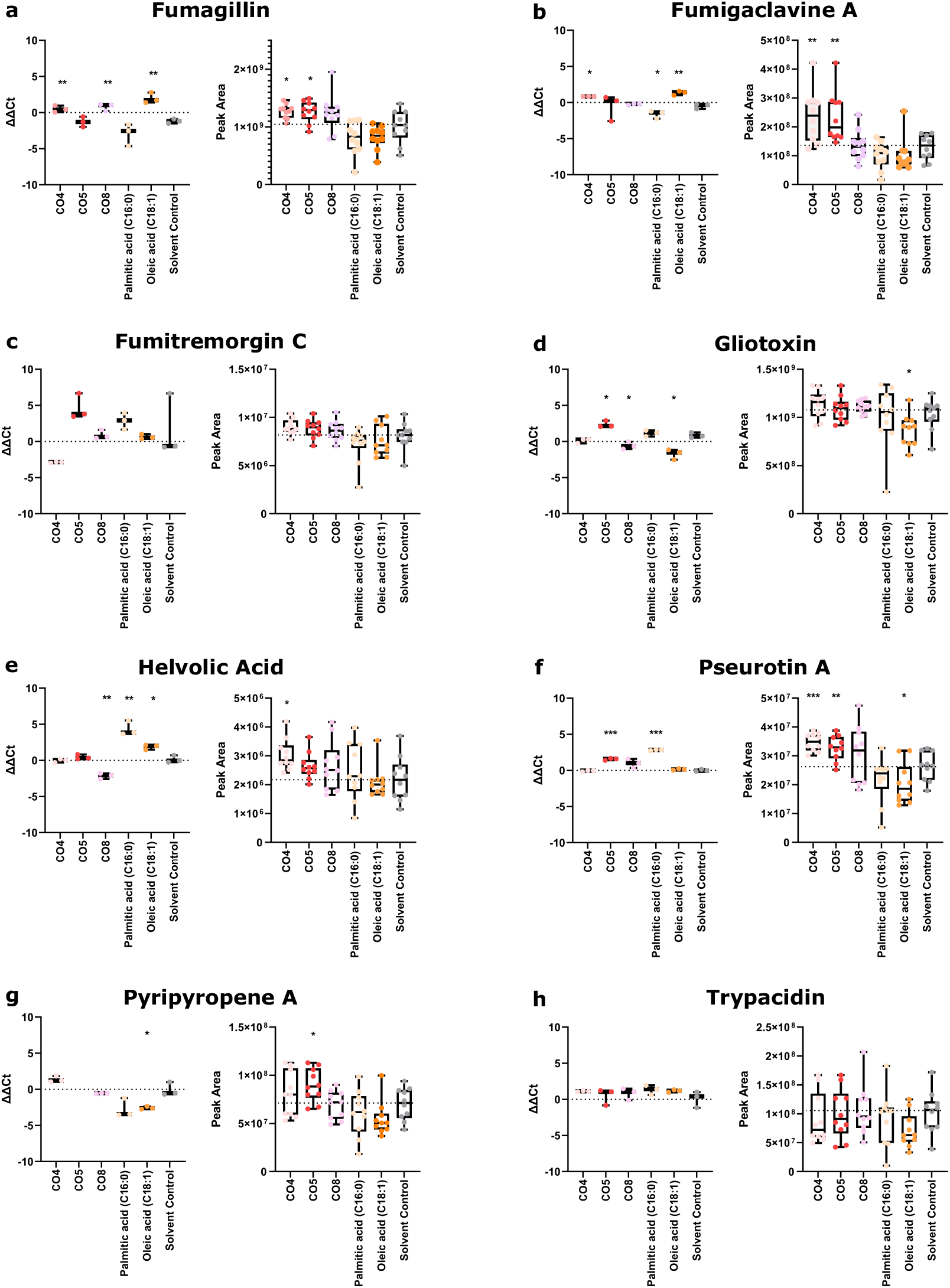
Metabolic gene expressions and peak area for confirmed specialized metabolites. Welch’s ANOVA p-values were (a) < 0.0001 for ΔΔCT data and 0.0054 for peak area, (b) 0.0004 for ΔΔCT and < 0.0001 for peak area, (c) 0.0002 for ΔΔCT and 0.0385 for peak area, (d) < 0.0001 for ΔΔCT and 0.0081 for peak area, (e) 0.0006 for ΔΔCT and 0.0230 for peak area, (f) < 0.0001 for ΔΔCT and peak area (g) 0.0005 for ΔΔCT and 0.0054 for peak area, and (h) was not significant. The gene responsible for pyripyropene A production was not expressed under CO5 treatment. Unpaired Welch’s t-test was used to compare individual treatments to the solvent control resulting in (*) indicates p-value < 0.005; (**) indicates p-value < 0.01; (***) indicates p-value < 0.001; and (****) indicates p-value < 0.0001.

#### Revealing the dominant compounds and highly influenced known and putative metabolites - the direct route

The influence of chitooligosaccharides and lipids on the production of known and putative metabolites by *Aspergillus fumigatus* at 25°C was analyzed using the direct route as shown in Figs. 4 a-c. The bipartite networks provide a visual representation and enable a clear distinction between the effects of these treatments, as shown in Fig. 4a. While chitooligosaccharides resulted in an up-regulation of the identified metabolites, the lipids showed a down-regulation. The network centrality measure of node out-strength of the treatments reveals that CO4 has the highest effect on triggering metabolite production, followed by CO5, then oleic acid. The treatments CO8 and palmitic acid showed a minor influence on inducing metabolite production. This was expected as those two treatments influence the production of only one metabolite with low values of log_2_ fold change. Moreover, the putative metabolite nidulanin A possesses the highest node in-strength amongst the metabolites as it is the most regulated metabolite, influenced by the chitooligosaccharides CO4 and CO5. These results align with the inferences drawn from the UpSet and volcano plots in Figs. 2 a-f.

**Figure 4:**
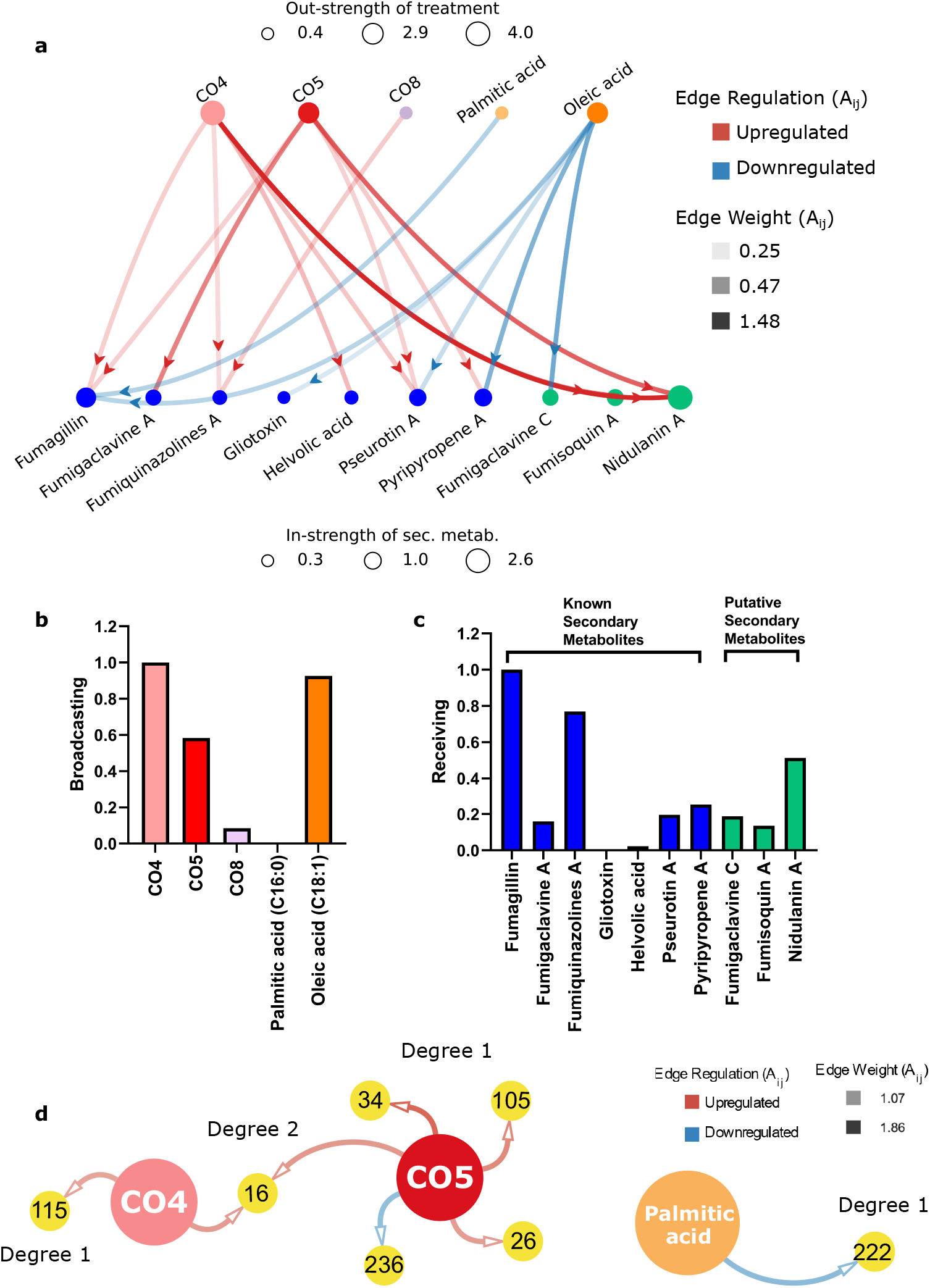
Network analysis-based direct and auxiliary routes for revealing the relationship between treatments and metabolite production in *Aspergillus fumigatus* at 25°C. (a) Bipartite network of treatments and known and putative metabolites. The nodes representing the metabolites are classified and colored coded for known and putative metabolites (blue and green, respectively). The transparency and color (red or blue) of the edges represent the log_2_ fold change and up- or down-regulation of the metabolites, respectively. The sizes of the nodes denote the network centrality measure of node strength. (b) Network centrality measure of PageRank of the treatments (broadcasting PageRank values). (c) PageRank measures the known and putative metabolites (receiving PageRank values). (d) Auxiliary route to assess the production of unknown analytes. A bipartite network of all analytes for their treatments (analyte IDs correspond to tables in S1 File providing mass to charge ratios (m/z), retention times, linear fold change, log_2_ fold change, p values, and f values). Degrees indicate the production of an unknown analyte by a single or multiple treatment(s) applied separately. Degree 1 are analytes induced by one treatment; Degree 2 are analytes produced by two different treatments. The weights and colors (red and blue) of the edges illustrate the log_2_ fold change of up- and down-regulation triggered by the treatments compared to the solvent control.

The network centrality measure of PageRank considers various factors, such as the number of edges from or to a node and the relative importance of nodes based on their connections to highly and uniquely connected nodes, to determine the most influential nodes in a network. The PageRank measure has been used extensively in various metabolic network analysis [3]. Due to the nature of metabolic interactions, variations in the PageRank measure has also been introduced [19]. We provide details on the differences between the node strength and PageRank measures in **Materials and methods** section. For the treatments, the ability to be influential at triggering metabolite production is measured by the broadcasting version of the PageRank measure. In contrast, the ability of metabolites to be receptive to treatments is denoted by the receiving version of the PageRank measure, as shown in Figs. 4b and 4c, respectively. The results, which are minimum-maximum normalized (values between 0 and 1, denoted herein by normalized broadcasting PageRank 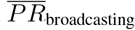 and normalized receiving PageRank 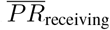), indicate that CO4 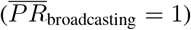 is the most effective treatment, followed by oleic acid 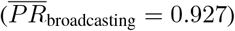. The bipartite network shows that CO4 triggers six metabolites compared to five triggered by oleic acid. Both treatments trigger two unique metabolites (helvolic acid and fumisoquin A by CO4; gliotoxin and fumigaclavine C by oleic acid). Nonetheless, CO4 has a higher influence on triggering the production of a unique metabolite, fumisoquin A. The broad number of metabolites activated by the CO4 treatment with a more prominent effect shows the wider impact of CO4 on triggering metabolites.

The current modeling framework reveals oleic acid to have a high impact on the production of metabolites even at 25°C. Oleic acid 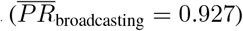 has a higher broadcasting PageRank value than CO5 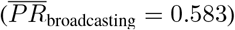, contrary to the node out-strength values (CO5 has higher out-strength than oleic acid as shown in Fig. 4b). The higher broadcasting PageRank value of oleic acid is attributed to its ability to uniquely trigger two metabolites (gliotoxin and fumigaclavine C) compared to just one by CO5 (fumigaclavine A). Furthermore, palmitic acid 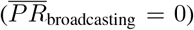 has the least broadcasting PageRank measure followed by CO8 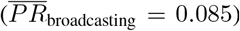, contrary to the results of node out-strength (CO8 has the least out-strength as shown in Fig. 4a). This change results from palmitic acid being connected to a highly receptive metabolite, fumagillin, triggered by many treatments. Thus, palmitic acid is not a unique treatment. CO8 is related to a unique metabolite, fumiquinazolines A, which is not triggered by many treatments.

The receiving PageRank measures of the metabolites reveal that the known metabolite fumagillin (normalized receiving PageRank, 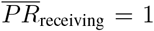) followed by fumiquinazolines A 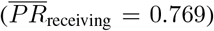 have much higher receptivity at being triggered by treatments compared to the putative metabolite nidulanin A 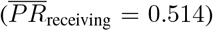. This result is contrary to that provided by the node in-strength, which showed nidulanin A to be the most influenced metabolite (Fig. 4a). While nidulanin A is the most regulated metabolite, its production was only activated by CO4 and CO5 treatments, producing many other metabolites. Therefore, the uniqueness of nidulanin A for being triggered is reduced. Furthermore, even though fumiquinazolines A has a much lower node in-strength value than nidulanin A and is only activated by CO4 and CO8, the latter treatment uniquely triggers this metabolite. This unique relationship with fumiquinazolines A is also one of the reasons why CO8 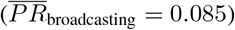 has a higher broadcasting PageRank value than palmitic acid 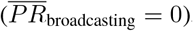, as discussed above.

These observations shown with the direct route could not be inferred using traditional methods like UpSet or volcano plots. Also, since the gene cluster for nidulanin A has been identified in all *Aspergillus* spp. and yet it has not been described in *A. fumigatus* [6, 52], CO4 and CO5 could be used as treatments for the characterization of this metabolite in *A. fumigatus*. Lastly, many of these known and putative metabolite peaks might still fall into a peak noise. Although a peak cutoff was initially used in MAVEN to identify bona fide peaks, the auxiliary route is used to identify known and unknown analytes or metabolites highly produced in response to a particular treatment using an untargeted metabolomics approach.

#### Revealing the dominant compound and highly influenced unknown analytes - the auxiliary route

The auxiliary route follows an untargeted metabolomic profiling of the treatments. The auxiliary route illustrated in Fig. 4d demonstrates our ability to potentially isolate highly produced known and unknown analytes that exhibit a log_2_ fold change greater than 1 for future experimentation and characterization. In the direct route, we had a peak area cutoff of 5× 10^5^ to detect signals with significant peaks between a treatment and solvent control. We hence allowed the identification and confirmation of known metabolites. Although our peak area cutoff of 5 × 10^5^ eliminates most noise, it is a linear cutoff and leaves some residual noise in the analysis due to column creep. Therefore, we first curated the dataset for the auxiliary route from the experimental study using baseline correction preprocessing tools. Fig. 17 shows the elimination of column creep, creating a new peak area cutoff suggested by Trevjño et al. [59]. In order to ensure the smallest amount of noise within our data, peak picking for our untargeted approach was done with GridMass [59] peak detection using a conservative threshold for peaks such that no peaks under 3 × 10^7^ would be considered. Peaks were aligned using RANSAC alignment. Peak data were then matched to known profiles in KEGG and LipidMap [29, 27, 28, 18]. Peak significance was determined upon FCROS scoring [16]. Non-significant analytes were not included in the network. An interactive map of the network illustrating the details of each analyte (m/z ratio, retention times, p-values, etc.) is available at https://web.eecs.utk.edu/~mlane42/. The **Materials and methods** section provides further information about the data curation for building the bipartite networks for the auxiliary route.

The untargeted extraction of statistically relevant peaks using the auxiliary route can yield a significant number of analytes for potential exploration. The edges and neighbors of the nodes in the network can be used to determine which analytes to be first considered for targeted exploration. Analytes of particular interest express both regulation and control depending on the treatment considered. Additionally, analytes of extreme up- and down-regulation can be of interest along with the node degree values of the analytes.

A network of all significant peaks (i.e. all significant metabolites regardless of log_2_ fold change intensity) can be found in **S1 Appendix** (Fig. 18). Fig. 4d only illustrates the significant peaks with log_2_ fold change greater than 1. The total number of interactions (edges in the network) found with the auxiliary route is considerably higher compared to the network in the direct route. Thus, the method reveals more information regarding the interactions amongst the treatments and metabolites than the direct route. Interesting artifacts are revealed from the network built using data at 25°C. At 25°C, analytes ID 26 and ID 105 were matched to fraxetin via the KEGG database query. These were found to be upregulated with respect to the CO5 treatment. Analyte ID 222 was matched to ICAS#18 from the LipidMap database. Palmitic acid significantly down regulates ICAS#18 (ID 222) with respect to the control. At the same time, no single highly produced analyte is triggered by all three chitooligosaccharides. Analyte 16, however, is significantly produced by both CO4 and CO5 treatments.

All analytes of log_2_ fold change intensity greater than 1 except for analyte ID 16 are of degree 1. When not taking log_2_ fold change intensity into account, there exist 41 analytes of degree 1, 25 analytes of degree 2, 8 analytes of degree 3, 3 analytes of degree 4, and 1 analyte of degree 5 (see **S1 Appendix** and Fig. 18). These findings suggest that although treatments can commonly start the production of the analytes considered, the treatments used in the current study have a higher tendency to uniquely trigger analytes, which agrees with the UpSet and volcano plots and direct route analysis.

Analytes that immediately spark interest are the four unknown analytes with log_2_ fold change intensity greater than 1 with IDs 16, 34, 115, and 236. Additional analytes of interest are those with opposing log_2_ fold changes between treatments, whereas all remaining analytes within the networks have aligned log_2_ fold changes. Though we do not see analytes with opposing log_2_ fold changes in the network with all log_2_ fold changes greater than 1 (Fig. 4d), when considering all edges we do see analytes with opposing log_2_ fold change intensities with IDs 13, 35, 115, and 188 (Fig. 18). A detailed discussion of the findings from the full data can be found in **S1 Appendix**.

### Data-driven validation of direct and auxiliary routes with treated samples at 37°C

#### Analyte and metabolomic production induced by treatments

We analyzed the LC/MS data with treated samples grown at 37°C, which revealed that all individually applied treatments significantly induce the production of analytes compared to the solvent control, as shown by the UpSet plot in Fig. 5a. No treatments were co-applied to the fungus. As expected, the total number of analytes is lower at 37°C than at 25°C. In total, 1,807 significant analytes were detected (Fig. 5a). The percentage of unique analytes produced by CO4 was 25.6%, by CO5 was 13.6%, by CO8 was 17.2%, by palmitic acid was 10.6%, and by oleic acid was 5.0%. These results indicate that at 37°C, both short-chain and long-chain COs influence specific metabolomic pathways, unlike at 25°C. At 37°C, the CO8 treatment induced the production of particular metabolites by the fungus better than at 25°C. Contrary to what was observed at 25°C, oleic acid showed less influence on analyte production at 37°C compared to palmitic acid. It is also worth noting that most analytes were uniquely produced by a treatment rather than shared by multiple treatments at 37°C.

**Figure 5:**
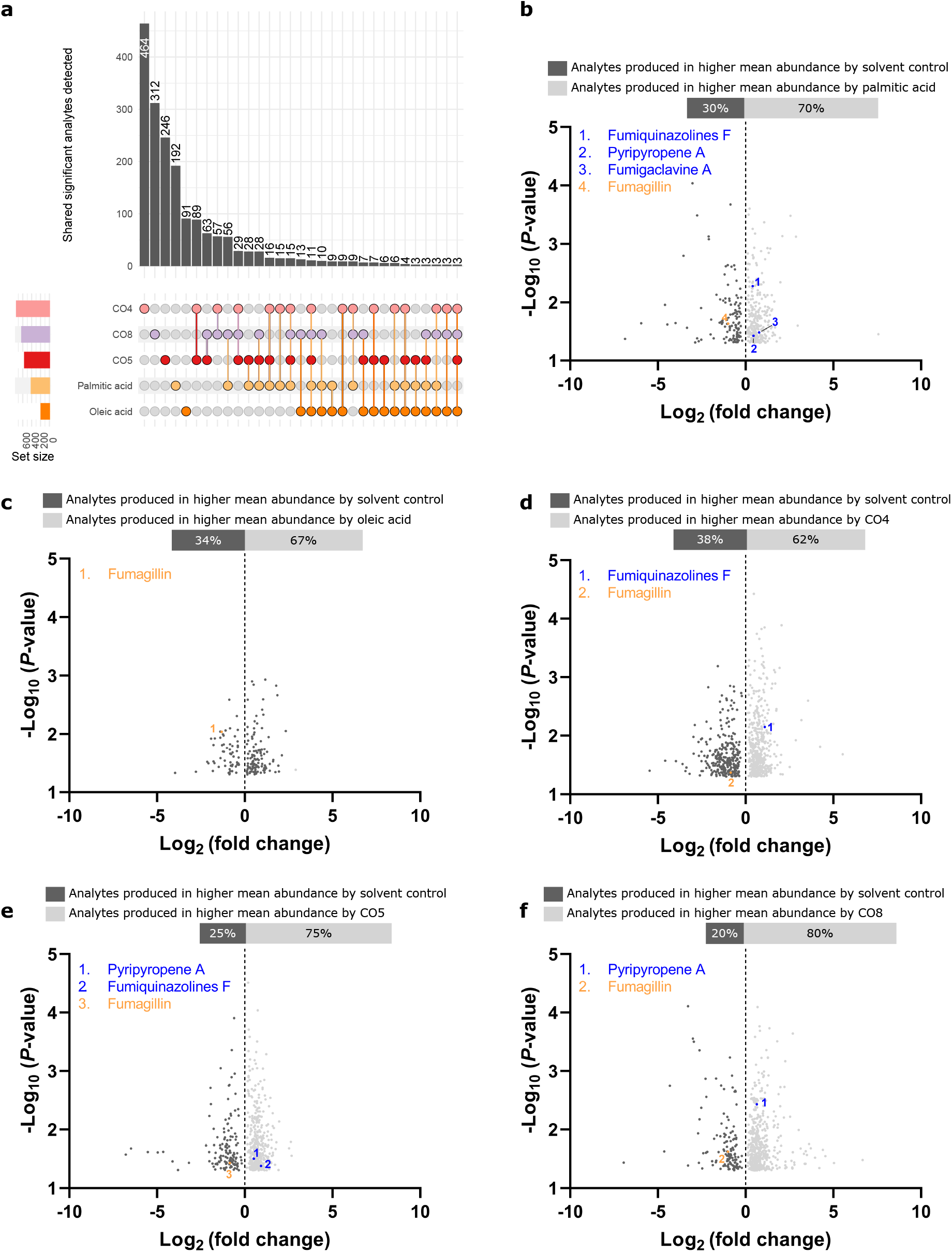
Metabolomic outputs of *Aspergillus fumigatus* at 37°C induced by lipid and chitooligosaccharide treatments. (a) UpSet plot denoting the number of significant analytes produced by individually applied treatments. Multiple treatments induced some analytes. (b-f) Volcano plots identifying the known and putative metabolites and unknown analytes triggered by (b) palmitic acid, (c) oleic acid, (d) CO4, (e) CO5, and (f) CO8 as compared to the solvent control.

With the LC/MS data produced at 37°C, we made volcano plots based on the log_2_ fold change and − log_10_ (p-values) between a treatment and the solvent control, as shown in Figs. 5 b-f. Although all treatments had a significant differential production of analytes compared to the solvent control at 37°C, it was less than that observed at 25°C, as expected (Figs. 2 b-f). COs had a higher mean of abundance ranging between 62% to 80% (Figs. 5 d-f), whereas lipids had between 67% to 70% (Figs. 5 b-c). These results mean that COs reduce the abundance of analytes at 37°C compared to 25°C. However, lipids showed a higher mean quantity of analytes at 37°C compared to 25°C. These changes in the results indicate that treatments influence the production of analytes, but environmental cues like temperature also have a significant effect. Therefore, further investigations to elucidate metabolites should be conducted at 25°C for the CO treatments and 37°C for lipids. Interestingly, nearly all treatments increased the production of the same known or putative metabolites, fumiquinazolines F, fumigaclavine B, or pyripyropene A, whereas all treatments reduced the production of fumagillin at 37°C. The regulation of the latter compound was different at 25°C, where the lipid treatments reduced its production and the short-chain COs improved it. This indicates that lipids might be compounds that downregulate fumagillin production constantly across temperatures and could be further investigated as therapeutic molecules.

#### Revealing the dominant compounds and highly influenced known and putative metabolites - the direct route

The results of the direct route to reveal the influence of the treatments on the production of specialized metabolites in *Aspergillus fumigatus* at 37°C are shown in Figs. 6 a-c. As expected, fewer analytes and known or putative specialized metabolites were produced at 37°C in both solvent controls and treatments, as shown in Fig. 6a and previously discussed using the UpSet plots in Fig. 5a. Furthermore, there is no clear distinction on how the two treatments regulate the production of metabolites. Both chitooligosaccharides and lipids showed a positive and negative impact on metabolite production at 37°C, whereas, at 25°C, chitooligosaccharides upregulated the production of metabolites and lipids down-regulate it (Fig. 4a). It is worth noting that the metabolites fumagillin and pyripyropene A are the only metabolites that were uniquely triggered at both temperatures; however, the treatments that started the production of these two metabolites were different depending on the temperature. The bipartite network representation further clarifies such pathways and regulations.

**Figure 6:**
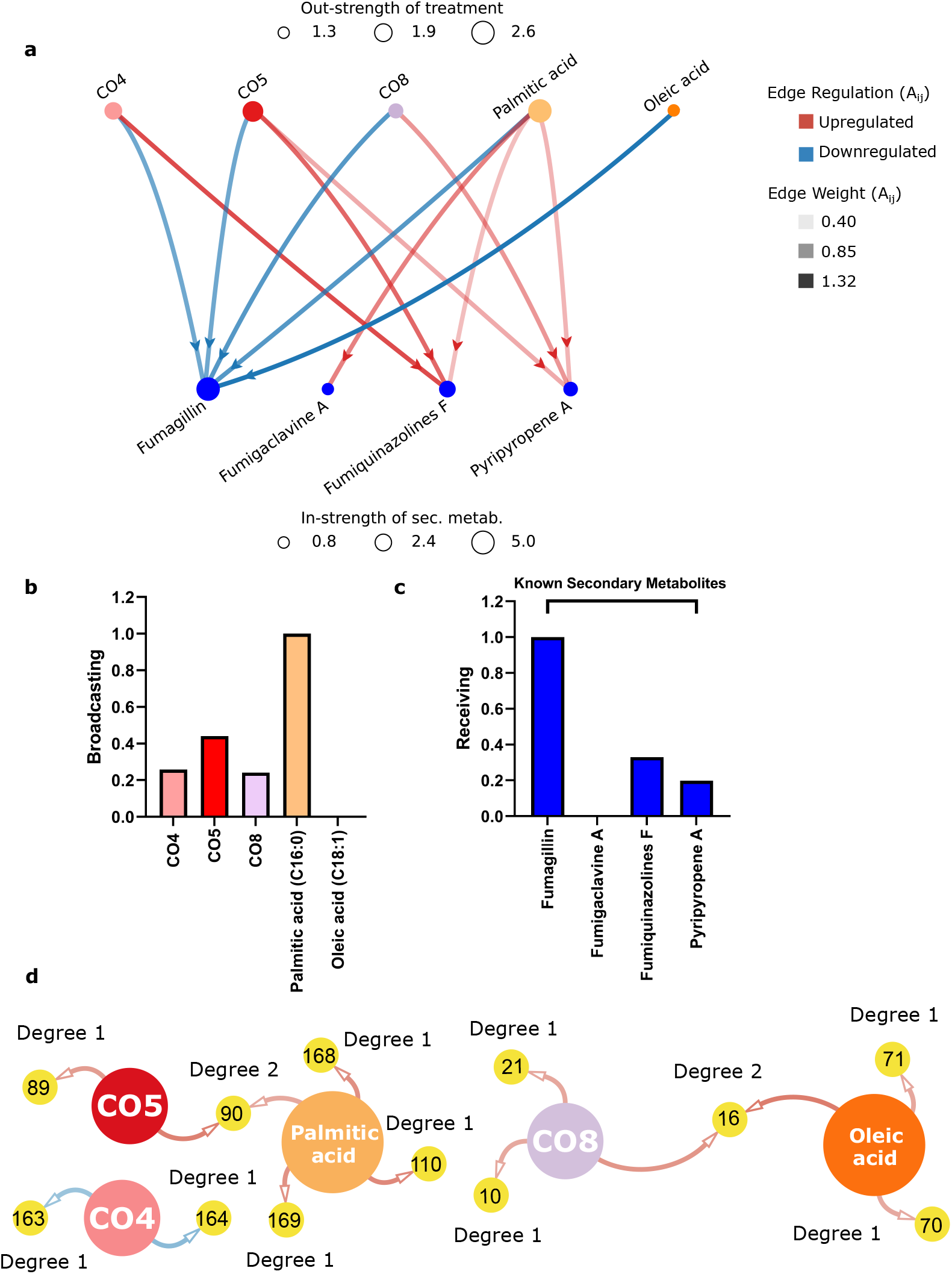
Network analysis-based direct and auxiliary routes for revealing the relationship between treatments and metabolite production in *Aspergillus fumigatus* at 37°C. (a) Bipartite network of treatments and known and putative metabolites. The nodes representing the metabolites are classified and colored coded for known and putative metabolites (blue and green, respectively). The transparency and color (red or blue) of the edges represent the log_2_ fold change and up- or down-regulation of the metabolites, respectively. The sizes of the nodes denote the network centrality measure of node strength. (b) Network centrality measure of PageRank of the treatments (broadcasting PageRank values). (c) PageRank measures the known and putative metabolites (receiving PageRank values). (d) Auxiliary route to assess the production of unknown analytes. A bipartite network of all analytes for their treatments (analyte IDs correspond to tables in S2 File providing m/z values, retention times, linear fold change, log_2_ fold change, p values, and f values). Degrees indicate the production of an unknown analyte by a single or multiple treatment(s) applied separately. Degree 1 are analytes induced by one treatment; Degree 2 are analytes produced by two different treatments. The weights and colors (red and blue) of the edges illustrate significant up- and down-regulation triggered by the treatments compared to the solvent control.

With the limited number of data points, both the node strength and PageRank measures give similar results for identifying the effective treatments and most receptive metabolites, as shown in Figs. 6 b-c. The lipid palmitic acid 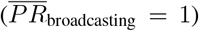 is the most effective treatment on metabolite production as it resulted in both up- and down- regulation of all metabolites analyzed. At 37°C, CO4 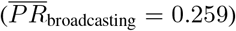 and oleic acid 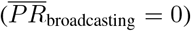 are poor at triggering the production of metabolites, contrary to what was observed at 25°C, where these treatments showed the most influence (Fig. 4b). The specialized metabolite fumagillin 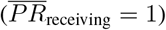 has the highest receptivity at being triggered by treatments, as shown in Fig. 6c. The same result was observed at 25°C.

#### Revealing the dominant compounds and highly influenced analytes - the auxiliary route

Considering the analytes from the network at 37°C shown in Fig. 6d, twelve significant analyte peaks with log_2_ fold change intensities greater than 1 were extracted from the processed data. All three CO treatments at 37°C triggered no shared analytes. Nor did the two lipid treatments trigger a shared analyte. Ten of the twelve analytes with log_2_ fold change intensities greater than 1 are uniquely started by a single treatment (i.e. analytes with node degree 1), as shown in Fig. 6d. Of the twelve significant extracted analyte peaks, eleven were identified from database queries. Analytes identified by KEGG were hellebrigenin 3-acetate (ID 10), fraxetin (ID 16), beta-cyclopiazonate (ID 70), phenylbutazone (ID 71), borrerine (ID 89), clofibrate (ID 110), alangimarine (IDs 163 and 164), and sulindac (IDs 168 and 169). LipidMaps identified 6’-hydroxysiphonaxanthin decenoate (ID 21). The only unidentified analyte was analyte ID 90, upregulated in both CO5 and Palmitic Acid treatments.

When considering all analytes (not only those with log_2_ fold changes greater than 1) as shown in Fig. 19 in **S1 Appendix**, there exist five analytes with opposing log_2_ fold changes (analytes with IDs 21, 70, 163, 164, and 168) compared to the four analytes at 25°C. Analytes 163 and 168 are both of degree 3. Analyte 163 is upregulated by both palmitic acid and CO8, yet downregulated by CO4. Analyte 168 is upregulated by palmitic acid, yet downregulated by both CO4 and oleic acid.

Oleic acid was reported as an inducer of germination in *Aspergillus fumigatus* at 37°C [53]. To our knowledge, none of the known metabolites identified were previously linked to germination in *A. fumigatus*. Therefore, it is tempting to speculate if one of the highly up-regulated unknown analytes could be the culprit behind the increased germination of this fungus at 37°C, which can be the target for future experiments. A detailed discussion of the findings from the full data can be found in **S1 Appendix**.

## Discussion and Conclusion

The effects of compounds like chitin and lipids on microbial metabolomic profiles are not fully elucidated and remain challenging to interpret. This study provides a data-driven modeling framework using network analysis to dissect the connection between exogenous inputs - biological compounds like lipids and chitooligosaccharides - and the metabolomic outputs - putative metabolites and unknown analytes - in the opportunistic human pathogen *A. fumigatus*. Bipartite networks with two classifications of nodes are built. The network nodes represent the treatments and specialized metabolites under consideration. The edges connecting the nodes represent the magnitude of up- or down-regulation of the specialized metabolites triggered by the corresponding treatments. We provide two routes to characterize the production of the specialized metabolites: (1) the direct route for the production of known and putative metabolites and (2) the auxiliary route for the production of unknown analytes. Moreover, we use network centrality measures of node strength and PageRank to rank the treatments and specialized metabolites. The treatments are ranked based on their ability to trigger the production of various specialized metabolites. The specialized metabolites are ranked based on their ability to be influenced by multiple treatments.

The results of the direct route reveal that the chitooligosaccharides, CO4, followed by the lipid, oleic acid, are the most dominant treatments to trigger a broad range of known and putative metabolites at 25°C. However, at 37°C, palmitic acid is the most effective treatment for metabolite production. We have also found that fumagillin is the most receptive specialized metabolite in *A. fumigatus* triggered by various chitooligosaccharides and lipids at both temperatures. Without the use of network-based measures like PageRank, this vital information would not be revealed by solely analyzing the total magnitude of up- or down-regulation of a metabolite caused by treatments, which in the current analysis for *A. fumigatus* would have resulted in the putative metabolite nidulanin A being identified as the most receptive specialized metabolite to be triggered by chitooligosaccharides and lipids when the fungus is exposed to a higher temperature.

The results of the auxiliary route illustrate that there is a far greater number of analytes significantly produced than the current amounts of known and putative metabolites. The significant analytes extracted at temperatures 25°C and 37°C comport not only with known and putative specialized metabolites but also with the rates at which the triggering of specialized metabolites have been witnessed in the previously discussed direct route. Moreover, new relations among treatments and metabolites are revealed as all found peak signals are considered in the auxiliary route. Of the significant analytes produced, there is a tremendous overlap in the activation of analyte production between treatments. We additionally found that by thresholding edges to be greater than or equal to 1 log_2_ fold change, we were able to paint a far more digestible illustration of how these treatments begin to interact with the up- and down-regulation of the untargeted analytes. When considering all significant analytes, we see that within the 25°C network, the lipid treatments tend toward down-regulation whereas the chitooligosacharides tend to up-regulate analyte production. Conversely, within the 37°C networks, we see that all treatments tend toward up-regulation of analytes except for CO4, which has a relatively balanced amount of up- and down-regulated induced analytes.

In Table 1 we summarize and compare the capabilities of the current framework with that of other state-of-the-art metabolic network analysis tools. Note that while the referenced tools have a vast array of capabilities, we are only mentioning the ones that are relevant to the current analysis. The insights about the most effective treatments and most influenced specialized metabolites are valuable for (1) validating known specialized metabolites through applied exogenous treatments or environmental cues and (2) discovering new specialized metabolites from putative metabolites and unknown analytes by genetic knockouts to characterize their gene clusters as depicted in post-analysis applications (Fig. 1). Ultimately our goal was to track how a treatment will elucidate the production of secondary metabolites. It is widely known that most biosynthetic gene clusters are silent under standard culture conditions resulting in minimal production of secondary metabolites. Our study can help researchers determine how their treatments will improve the production and accumulation of natural products. Those results can be validated through mass spectrometry analysis and comparison to fragmentation patterns from published datasets or commercial standards and through transcriptomic analysis to assess their biosynthetic gene expressions. Further confirmation can be done through knockout experiments and functional validation of the targeted biosynthetic gene clusters in post-analysis applications, which is outside the scope of this manuscript.

**Table 1:**
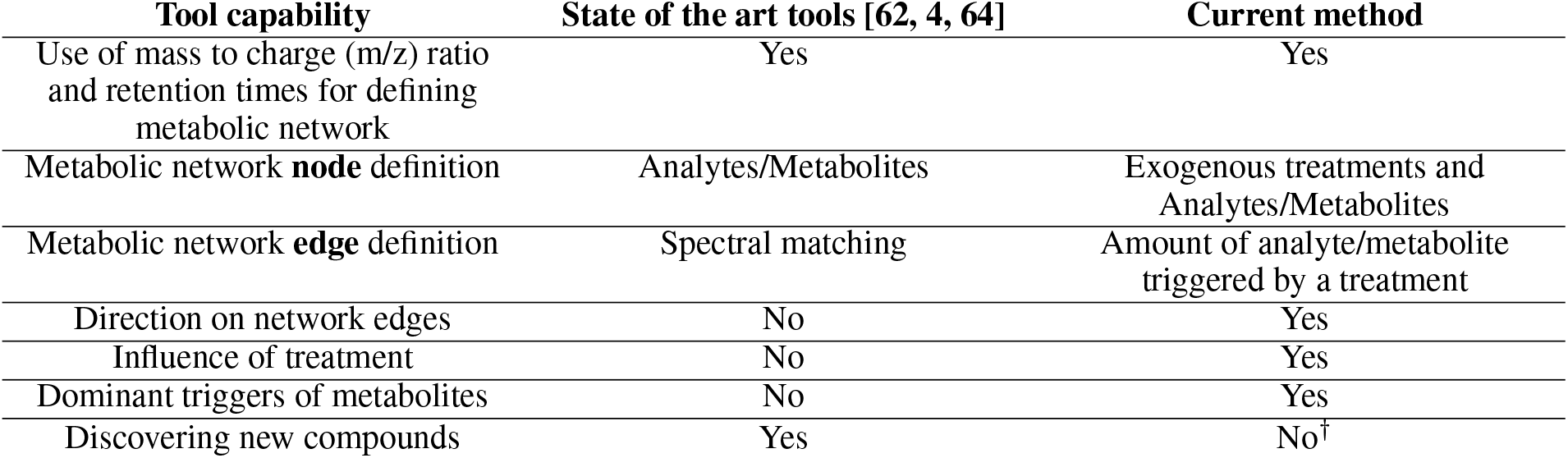
Comparison of relevant capabilities of state-of-the-art metabolic network analysis tools (GNPS [62], MetWork [4], and MetaboAnalyst [64]) with the current analysis. ^*†*^New compounds can be discovered by validating the identified important analytes through post-analysis applications elaborated in Fig. 1 - not performed in this study.

The cost of drug discovery can be exponential if experiments are designed by a trial-and-error method. We provide a potentially cost-effective solution through the direct and auxiliary routes (along with relevant codes on Link for researchers to use on publicly available or newly generated LC/MS datasets). The inferences obtained from our framework can be used as a guidance for industry partners and researchers to concentrate their efforts on natural product discovery and post-analysis applications for the most influential microbial metabolites and treatments.

## Materials and methods

### Network analysis

#### Bipartite network of treatments and specialized metabolites

A network (or graph), mathematically represented as 𝒢 (𝒱, ℰ, 𝒲),comprises a collection of vertices (or nodes) 𝒱 connected by edges ℰ representing the influence of the nodes onto each other [44]. The edges can be binary connections (0 or 1) or weights 𝒲). For our problem, we considered the treatments and specialized metabolites (known and putative metabolites and unknown analytes) as the nodes of a network, and the amount of change in the production of the specialized metabolites triggered by the treatments represents the weighted edges. Thus, the edge weights represent the magnitude of up- or down-regulation of the specialized metabolites affected by the treatments compared to a controlled case. Thus, the unidirectional edges representing the interaction of the treatments *j* on the specialized metabolites *i* can be represented by the elements of the adjacency matrix

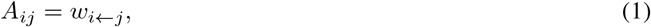

where *w*_*i←j*_ is the magnitude of change in concentration of *i* when exposed to treatment *j* measured in terms of the absolute value of the log_2_ fold change. Accordingly, the influence of specialized metabolites on treatments is zero, and the ***A*** matrix will have zero entries when *j* is a specialized metabolite. This results in a directed (unidirectional) network that can also be viewed as a bipartite network with the sources on one group and the specialized metabolites on the other, as shown in Figs. 4a and 6a. Moreover, red and blue colors can be assigned to the edges to depict the up- and down-regulation of the specialized metabolites by the treatments, respectively.

#### Network centrality measures

The network-theoretic measures of centrality are helpful to identify the most influential nodes in the network. We have found that the network centrality measures of node strength and PageRank [12] reveal physically informative information as we were able to validate some of the inferences from these measures with existing knowledge in literature. Moreover, these network centrality measures reveal additional information regarding important specialized metabolites and exogenous treatments. The node strength of node *i* is defined as the total interactions of the node *i*,

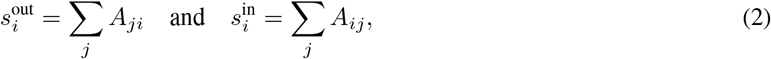

where 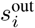 and 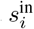 are the out- and instrength of *i*, respectively. According to the definition of the ***A*** matrix for our problem, out- and in-strengths were used to quantify the total influence of treatments for triggering the production of specialized metabolites and total receptivity of specialized metabolites to treatments, respectively (can be visualized by node size or circle radius as shown in Figs. 4a and 6a.

A limitation of the measure node strength is that it does not consider the number of connections from or to a node (the node degree). The number of metabolites triggered by treatment can show how diverse the effect of the treatment is. Similarly, the number of treatments affecting a metabolite denotes how easily they start it. Another criterion to understand the importance of a node is to quantify the relative importance of one node to another node to which the first one is connected. If a treatment triggers a metabolite that is also triggered by many other treatments, this emphasizes the ability of the metabolite to be influenced and not the unique ability of the treatment. The node strength measure does not highlight such factors.

The PageRank measure *x*_*i*_ of a node *i* is given by the normalized sum of centralities of its neighbors, defined by

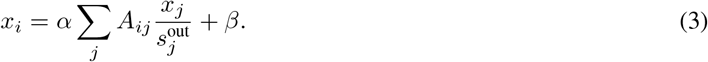

Here, *α* and *β* are positive constants used to balance between the normalized centralities of the neighbors and a minimum centrality value used for nodes that do not have connections. PageRank is the underlying method used by the Google search engine to rank a web page based on the number of directed links among the web pages, also considering the popularity of the web pages being linked to and from the web page. PageRank helps identify the relative importance of a node for the relevance of its neighbors. Moreover, it helps distribute the importance of high-interaction nodes to all the nodes’ connections and thus does not bias the centrality of the relationships. Therefore, if node *i* is connected to a well-connected node *j*, node *i* need not be an essential node just because it is pointed to by *j*. The node strength is overcome by the normalization, which distributes the strength of *j* to all its connections. For our problem, the directed version of the PageRank measure (not related to the “direct route”) was used to compute the broadcasting and receiving PageRank measures [23, 22] for the treatments and specialized metabolites, respectively. The importance of the treatments and specialized metabolites quantified by the strength (in- and out-strengths) and PageRank (broadcasting and receiving) are different and reveal important information regarding specialized metabolite production.

### Experimental design to elucidate metabolite production by treatments in *Aspergillus fumigatus*

#### Treatments

Short-chain COs - CO4 (IsoSep, product number 45/12-0050) and CO5 (IsoSep, product number 55/14-0050), and a long-chain CO - CO8 (IsoSep, product number 57/12-0001), were purchased from IsoSep (Tullinge, Sweden). The concentrations were 10^−8^ M in 0.005% EtOH/water, as described in [53]. Lipids - Palmitic acid, C16:0 (Millipore Sigma, item number P0500), and oleic acid, C18:1 (Millipore Sigma, item number O1008), were used at 10^−8^ M in 0.005% EtOH/water as described in [53].

#### Organism and inoculum

*Aspergillus fumigatus* strain Af293 was used in this study. The growth medium used was glucose minimal medium (GMM) broth. The Af293 strain was previously described in [50]. The medium was supplemented with various treatments short-chain or long-chain COs, and lipids at a final concentration of 10^−8^ M in the medium. The short-chain CO treatments were CO4 and CO5. The long-chain CO treatment was CO8. The lipids were palmitic acid (C16:0) and oleic acid (C18:1). The negative control for the analyses consisted of 0.005% EtOH, the solvent in which all the treatments were prepared. The spore concentration was adjusted to 10^6^ spores/ml of medium.

*Aspergillus fumigatus* spores were inoculated with exogenous CO or lipid standards at 25°C for six days and 37°C for four days. The solvent control was 0.005% EtOH. Liquid chromatography-mass spectrometry (LC/MS) of culture supernatants identified and quantified metabolites produced in response to different treatments. A standard analyte cutoff method evaluated with MAVEN v.8.1.27.32 [13] and XCMS v.3.7.1 [58] captured significant changes in analyte production across treatments and identified known specialized metabolites [52]. A stringent analyte cutoff method evaluated with mzMine2 [51] with column creep baseline corrections identified critical and specific mass to charge ratios and retention times of metabolites that are visualized using the bipartite networks. We provide additional details on the confirmation of the identities of the specialized metabolites in Section.

#### Metabolite profiling using ultra-high pressure liquid chromatography-mass spectrometry (UHPLC-MS) for the direct route

The effect of short-chain CO, long-chain CO, and lipid treatments on specialized metabolite production by *Aspergillus fumigatus* strain Af293 was assessed by UHPLC-MS analysis. About 10^6^ fresh spores were grown in 125 ml flasks containing 50 ml of GMM broth supplemented with the same treatments mentioned earlier. Two different growth conditions were assessed; the first consisted of incubation under 25°C and 250 rpm for six days, whereas the second consisted of incubation under 37°C and 250 rpm for four days. Optimal growth conditions for Af293 determine the difference in the time for growth at these conditions. After the incubation periods, fungal balls were collected and lyophilized to estimate the dry biomass. Dry biomass was not reported in this manuscript. For specialized metabolite analysis, 3 ml of supernatant were homogenized with 3 ml of ethyl acetate (Millipore Sigma, item number 270989). Organic and aqueous layers were separated by centrifugation at 3, 000 rpm for 5 min, and the organic layer was collected and dried. Samples were later resuspended in acetonitrile:water (50:50 v/v) and filtered through an Acrodisc syringe filter with nylon membrane (0.45*µ*m, Pall Corporation) into 1 ml HPLC vials. Samples were subjected to high-resolution UHPLC-MS analysis [50]. Data acquisition and processing for the UHPLC-MS were made using Thermo Scientific Xcalibur software version 4.2.47. Files were converted to .mzXML using MassMatrix MS Data File Conversion grouped by condition and run in MAVEN, an open-source software program [13] and XCMS open-source package (https://xcmsonline.scripps.edu/) using a pairwise comparison between treatments and controls with the parameter UPLC/Q-Exactive 3110 [58]. Volcano plots were based on analytes similar or uniquely produced by each treatment as determined by XCMS and illustrated in GraphPad Prism software version 9.0.0 (GraphPad, San Diego, California).

#### Feature extractions and differential expression analysis for discovery route

Raw mzXML chromatogram data were imported to mzMine2 [51]. Because of column creep, baseline correction was applied following protocols previously established in GridMass peak detection literature [59]. Baseline corrected data were cropped and preprocessed following procedures defined by [1]. Extracted peak features were quantile normalized by application and then log_2_ transformed; fold change rank ordering statistics were applied to all applications/controls to each combination of application and control to obtain fold change results [16]. Up- and down-regulated features were extracted with F values above 0.9 and below 0.1, respectively, with a p-value < 0.05. Parameter values can be viewed in Table 2.

**Table 2:** Preprocessing steps and parameter values for discovery route. The preprocessing steps and parameters used for postprocessing tools like GridMass, RANSAC, and FCROS to extract novel analyte peaks considering lower signal concentrations from the XCMS and MAVEN datasets.

|  |  |
| --- | --- |
| **Baseline Correction:** |  |
| MS Level | 1 |
| Use m/z bins checked m/z bin width | 1 |
| R engine | “Rcaller” |
| Correction Method | Asymmetric Baseline Corrector |
| Smoothing: | 10,000 |
| Asymmetry | 0.001 |
| **Peak Picking:** |  |
| Method | GridMass |
| Minimum Height | 3.00E+07 |
| m/z Tolerance | 0.01 |
| Minimum-maximum width time (min) | 0.03-0.08 |
| Smoothing time (min) | 0.1 |
| Smoothing m/z | 0.05 |
| False+: Intensity similarity ratio | 0.5 |
| False+: Ignore times | 0-0 |
| **Isotopic Peak Grouping:** |  |
| m/z tolerance | 0.001 m/z or 5.0 ppm |
| Retention time tolerance | 0.08 absolute (min) |
| Monotonic shape | unchecked |
| Maximum charge | 2 |
| Representative isotopes | “Lowest m/z” |
| **Alignment:** |  |
| Method | RANSAC Aligner |
| m/z tolerance | 0.1 m/z or 5.0 ppm |
| RT tolerance | 0.3 absolute (min) |
| RT tolerance after correction | 0.05 absolute (min) |
| RANSAC Iterations | 100 |
| Minimum number of points | 20.0% |
| Threshold value | 1.5 |
| Linear model | unchecked |
| Require same charge state | unchecked |
| **Duplicate Value Removal:** |  |
| Method | Duplicate Peak Filter |
| Filter mode | “New Average” |
| m/z tolerance | 0.001 m/z or 5.0 ppm |
| RT tolerance | 0.05 absolute (min) |
| **Gap Filling.** |  |
| Method | Same RT and m/z range gap filler |
| m/z tolerance | 0.001 m/z or 5.0 ppm |

#### Confirmation of specialized metabolite identities through fragmentation and comparison to standards

The identification of putative specialized metabolites was further interrogated via liquid chromatography tandem mass spectrometry analysis (LC-MS/MS) of specific m/z targets identified by MAVEN and XCMS software platforms. specialized metabolite identities were confirmed by MS/MS fragmentation pattern match through either in silico match to public databases and/or direct comparison to a commercially available standard (Figs. 7-16). Direct comparisons to standards offer the most confident identifications based on both MS/MS pattern match and confirmation of retention time (RT) by standard addition. Fragmentation patterns were matched to public online databases MASST (https://ccms-ucsd.github.io/GNPSDocumentation/masst/) and MassBank of North America (MoNA, https://mona.fiehnlab.ucdavis.edu/). Fumagillin, fumigaclavine A, fumitremorgin C, gliotoxin, helvolic acid, pseurotin A, pyripyropene A, and trypacidin were confirmed through both fragmentation pattern match and RT by comparison to a purchased standard. Fumiquinazoline A and fumiquinazoline F were confirmed through in silico fragmentation pattern match to public databases. Fumisoquin A, fumigaclavine C, and nidulanin A are classified as putative specialized metabolites because they could not be confirmed through fragmentation match to a database, nor were they available for purchase.

**Figure 7:**
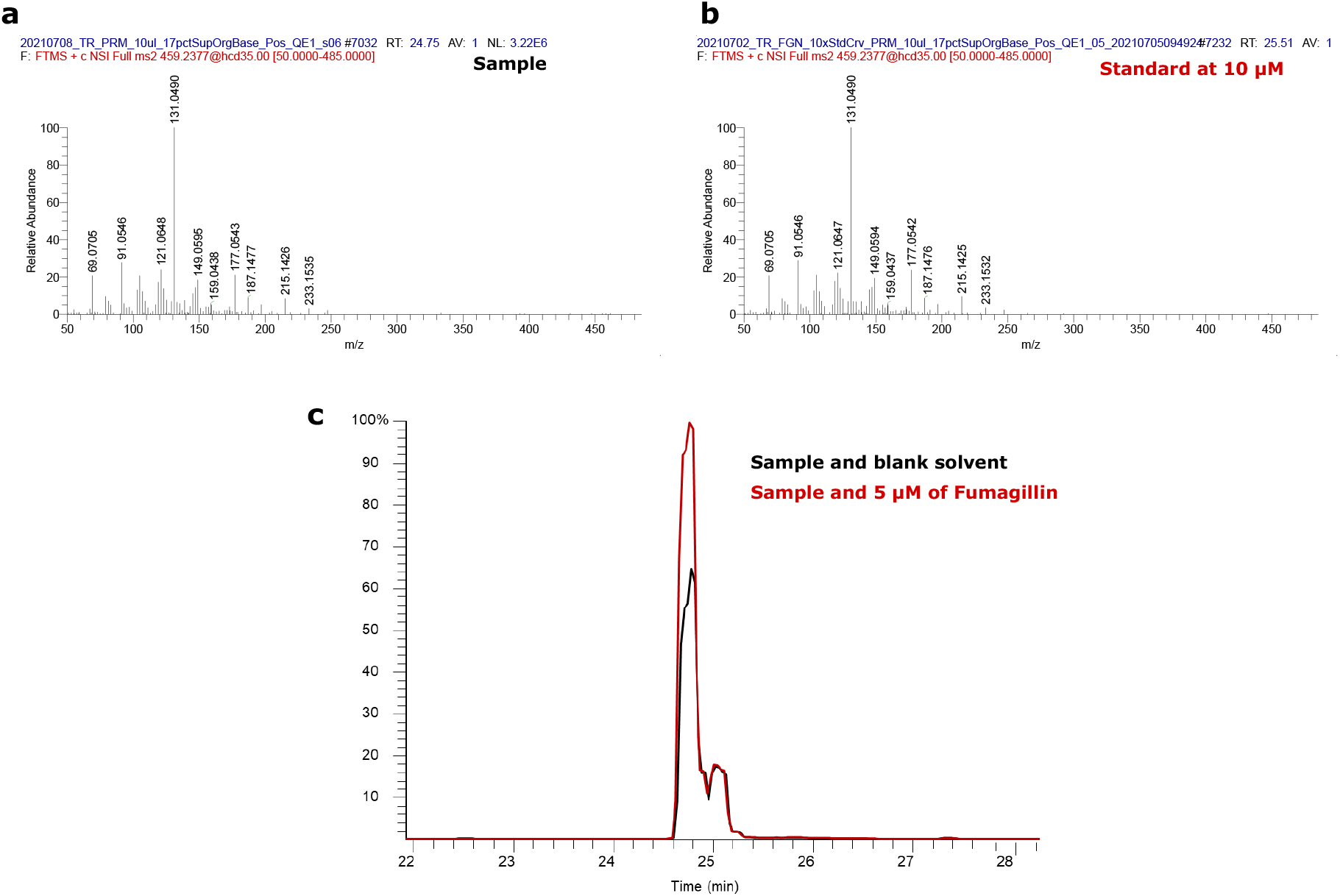
Fumagillin high resolution mass spectrometry dat. Fragmentation patterns of (a) a sample compared to (b) the standard, and (c) in silico spiking of the standard into the sample.

**Figure 8:**
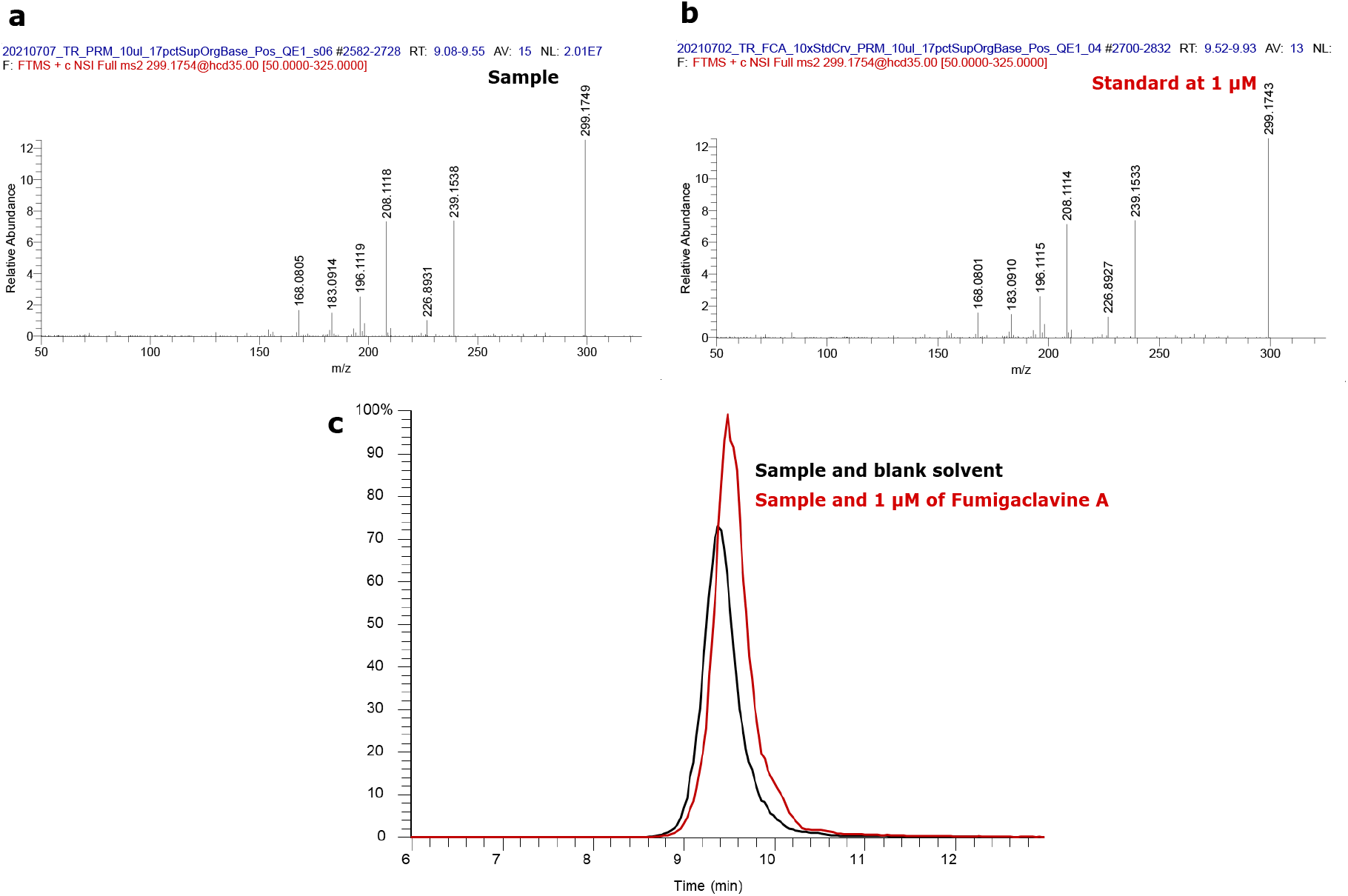
Fumigaclavine A high resolution mass spectrometry data. Fragmentation patterns of (a) a sample compared to (b) the standard, and (c) in silico spiking of the standard into the sample.

**Figure 9:**
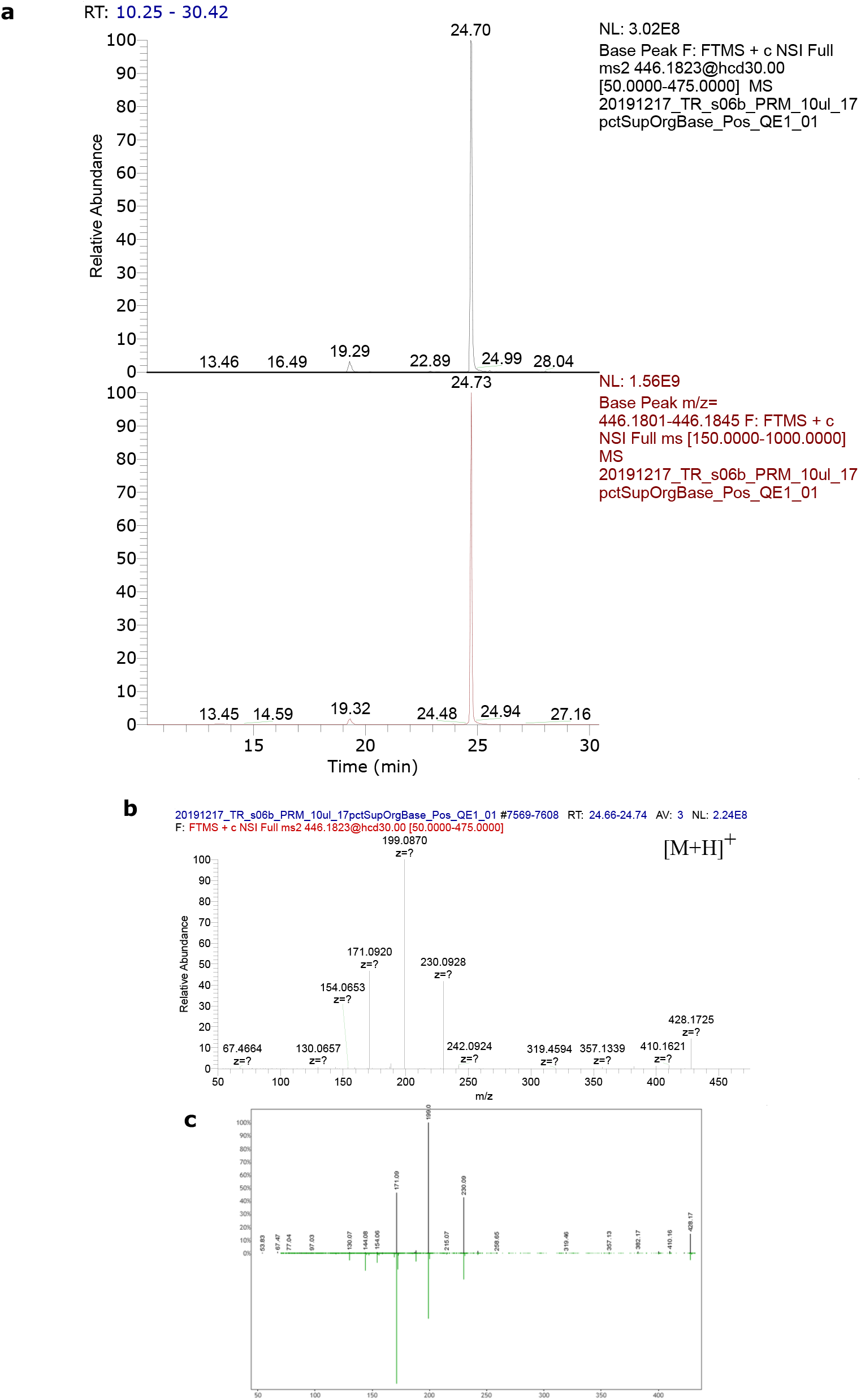
Fumiquinazoline A high resolution mass spectrometry data. (a) ion extract base peak from full mass spectrometry, (b) fragmentation mass spectrum in positive ionization mode, and c) associated profile with PubChem (CAS: 140715-85-1) and MASST.

**Figure 10:**
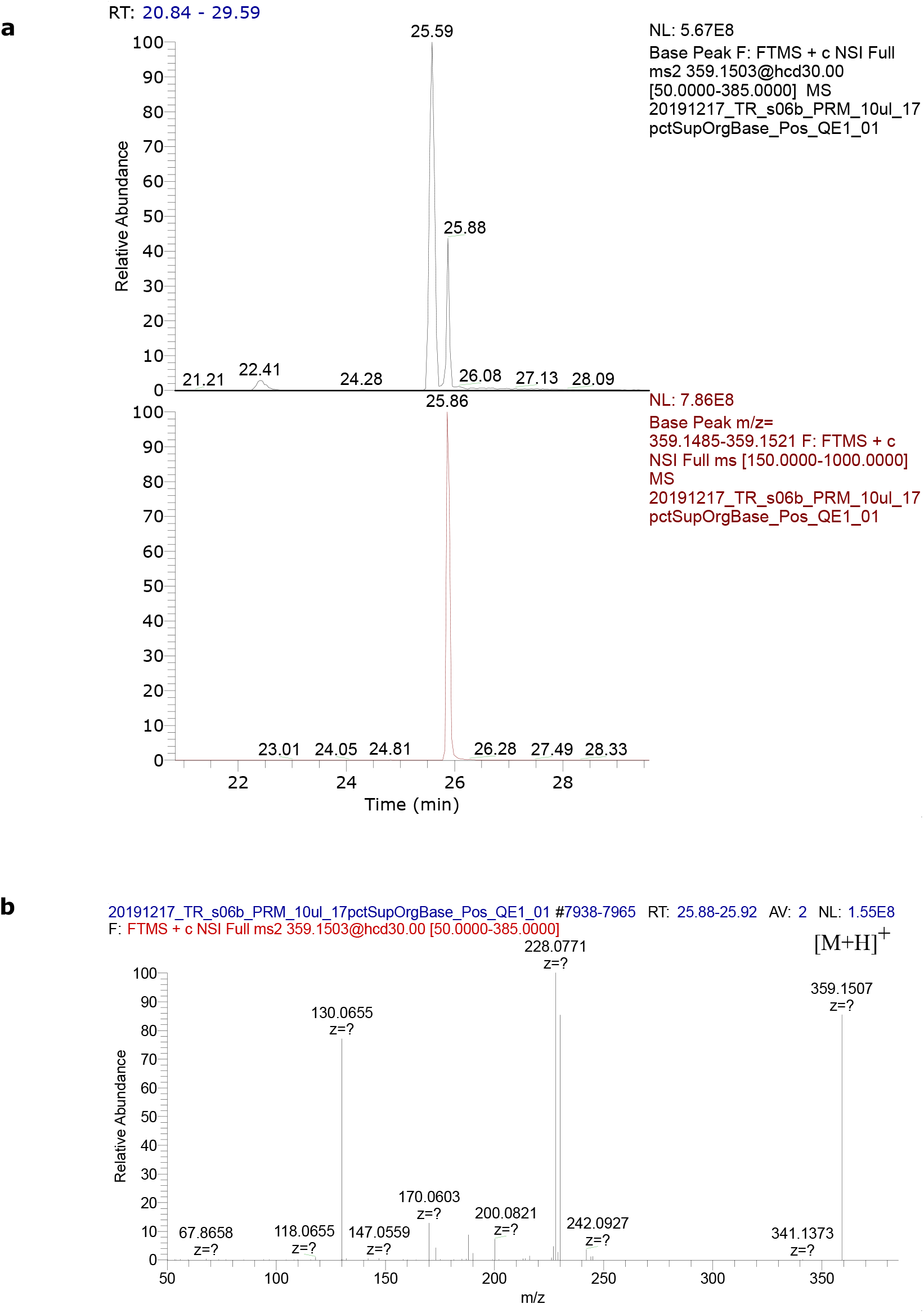
Fumiquinazoline F high resolution mass spectrometry data. (a) ion extract base peak from full mass spectrometry and (b) fragmentation mass spectrum in positive ionization mode. The associated profile was compared to MoNA (ID: AC00908 - https://mona.fiehnlab.ucdavis.edu/spectra/display/AC000908 and PubChem, CAS: 169626-35-1.

**Figure 11:**
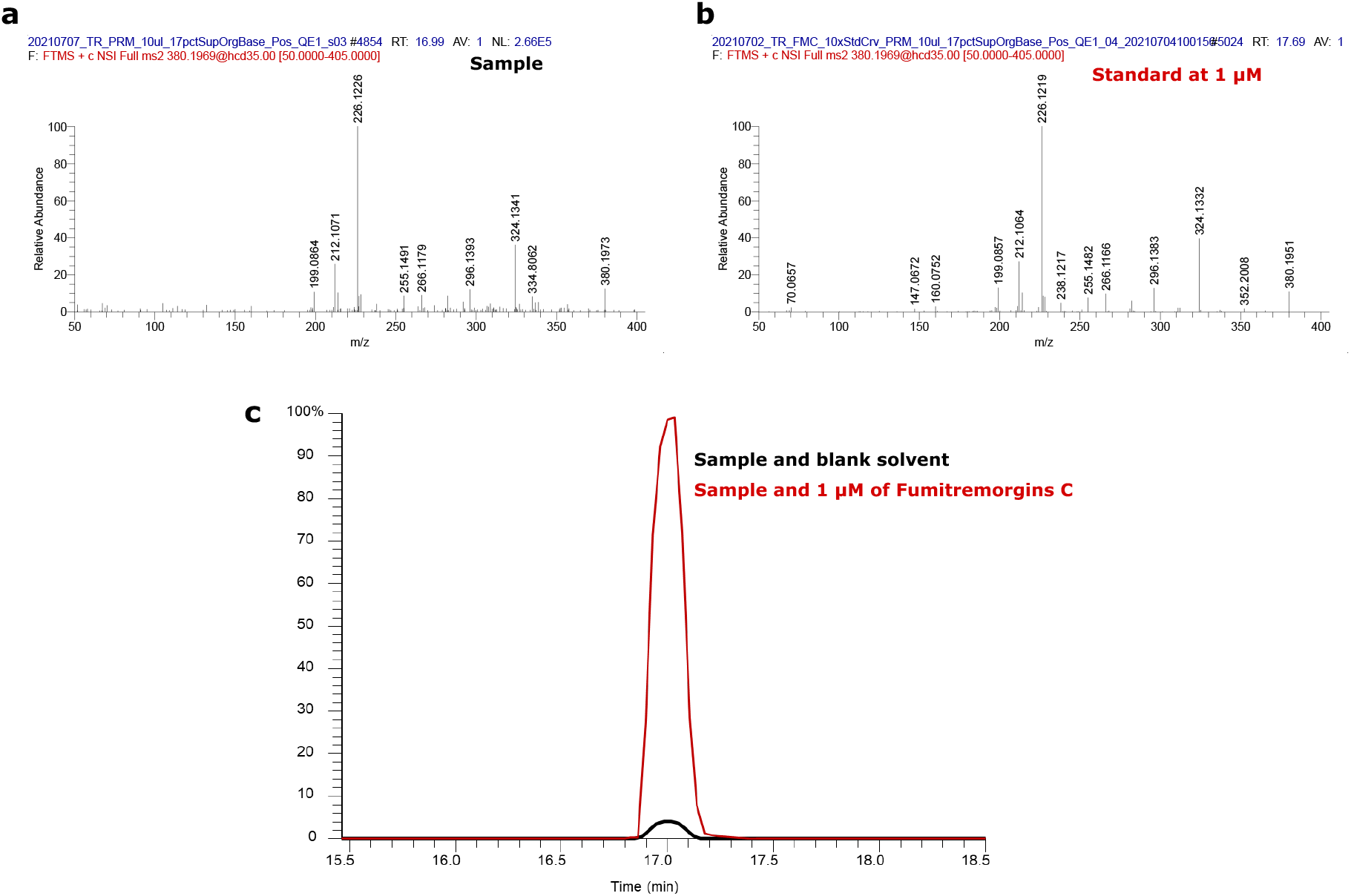
Fumitremorgin C high resolution mass spectrometry data. Fragmentation patterns of (a) the sample compared to (b) the standard, and (c) in silico spiking of the standard into the sample.

**Figure 12:**
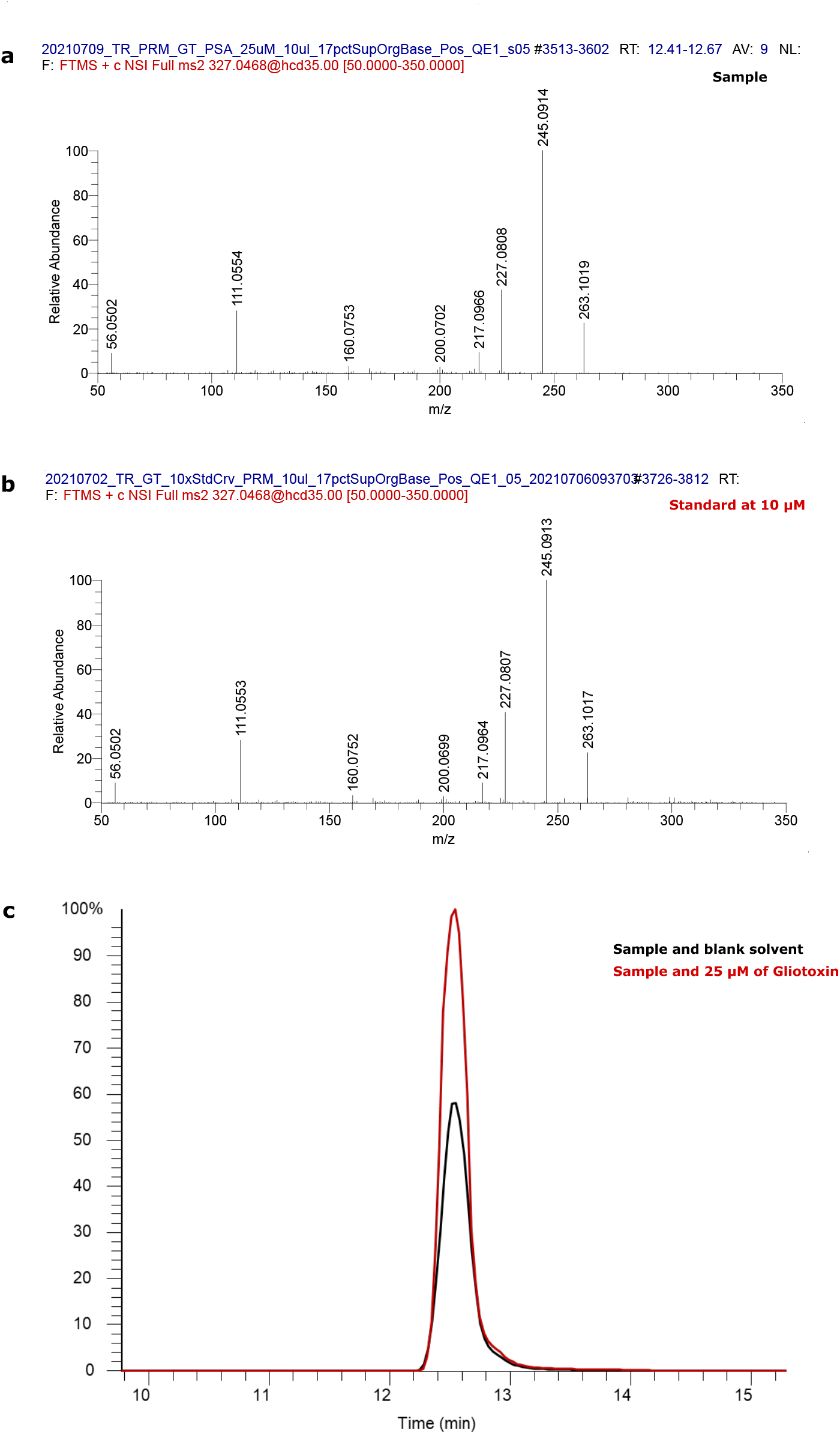
Gliotoxin high resolution mass spectrometry data. Fragmentation patterns of (a) a sample compared to (b) the standard, and (c) in silico spiking of the standard into the sample.

**Figure 13:**
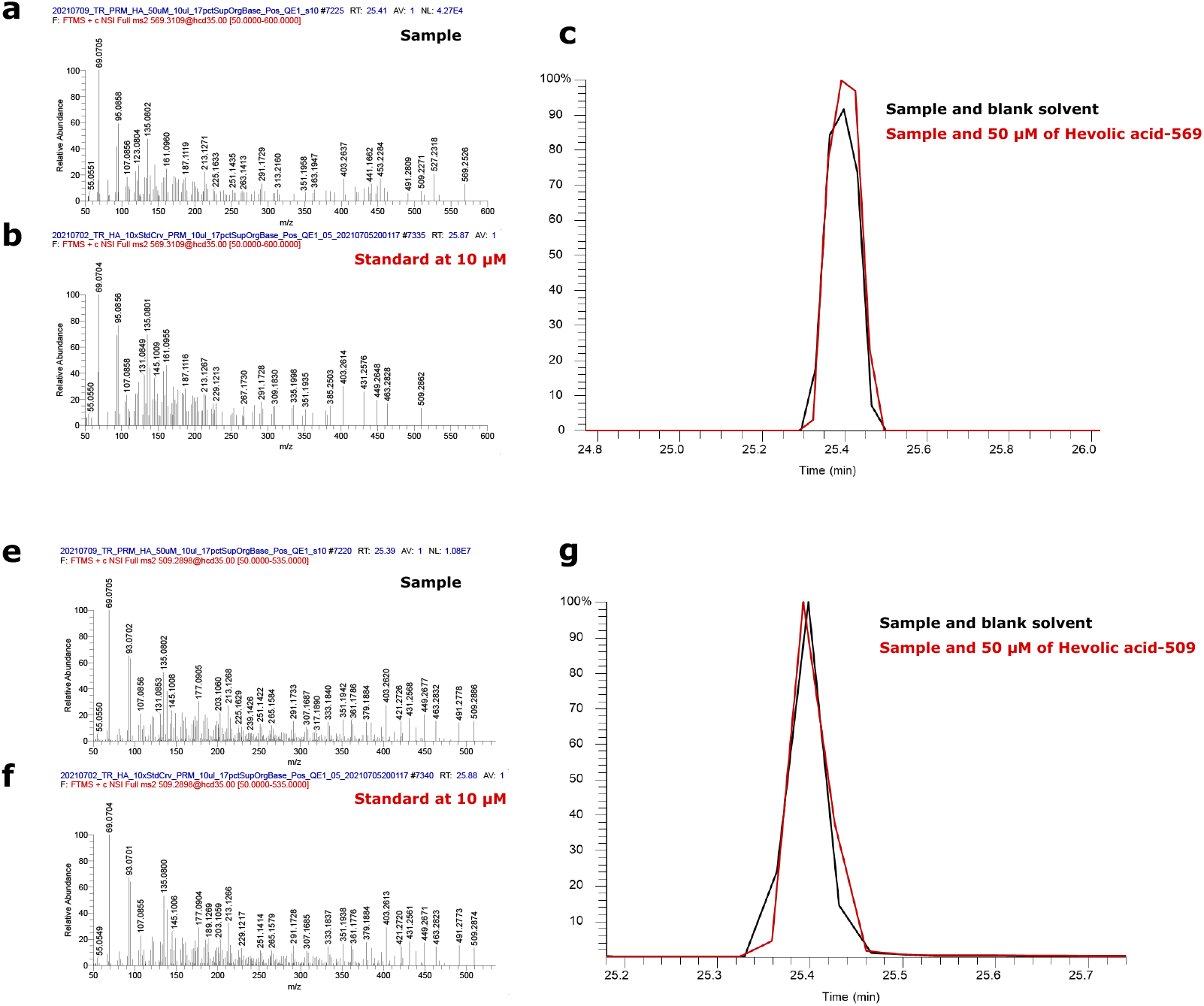
Helvolic acid high resolution mass spectrometry data. During ionization, a 569 ion will break apart and loses 60 Da to make a 509 ion. Therefore, both fragmentation patterns at 569 Da and 509 Da were analyzed. (a) sample compared to (b) the standard at 569 Da and (c) in silico spiking of the standard into the sample. (d) a sample compared to (e) the standard at 509 Da and (f) in silico spiking of the standard into the sample.

**Figure 14:**
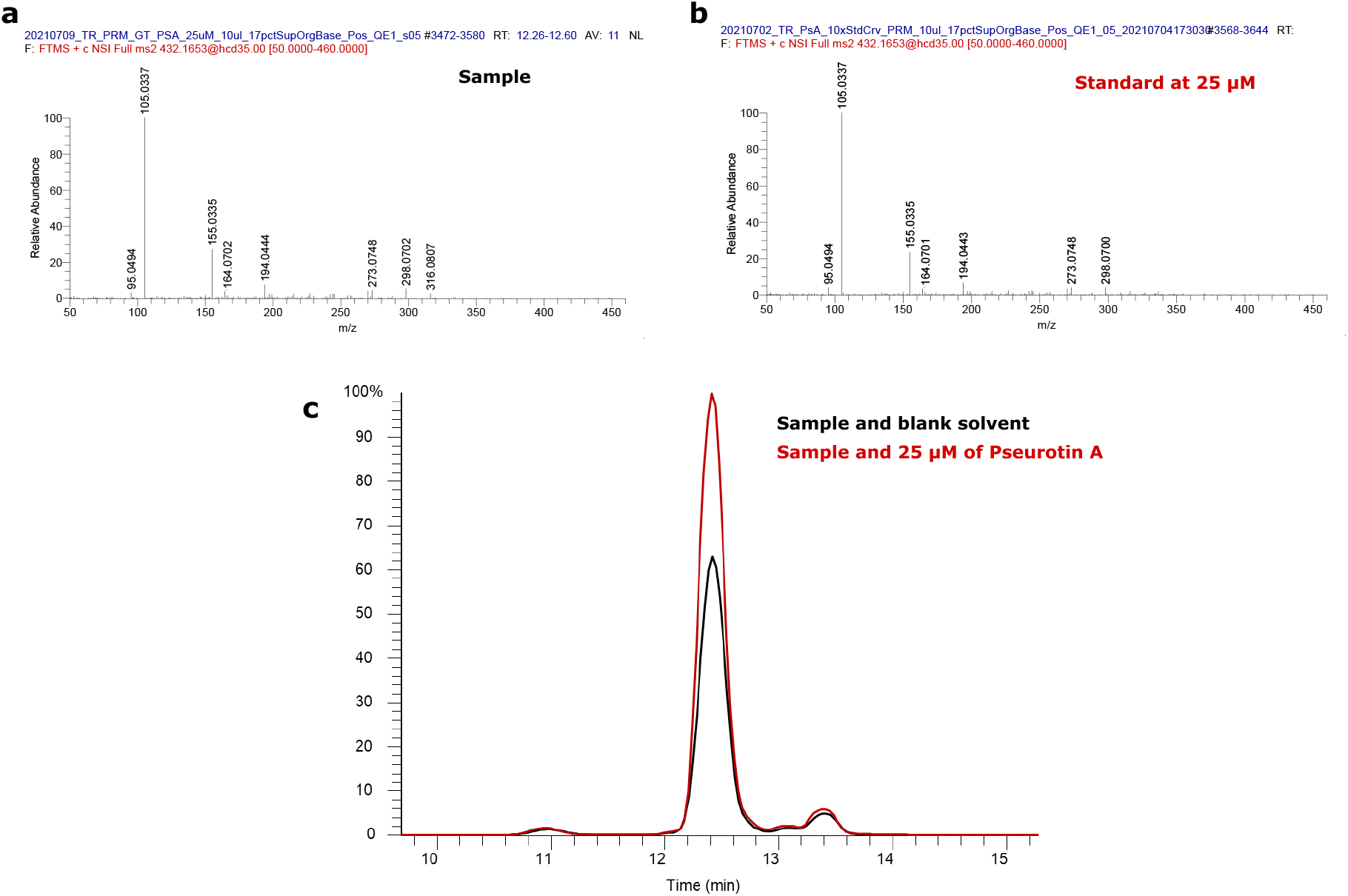
Pseurotin A high resolution mass spectrometry data. Fragmentation patterns of (a) a sample compared to (b) the standard, and (c) in silico spiking of the standard into the sample.

**Figure 15:**
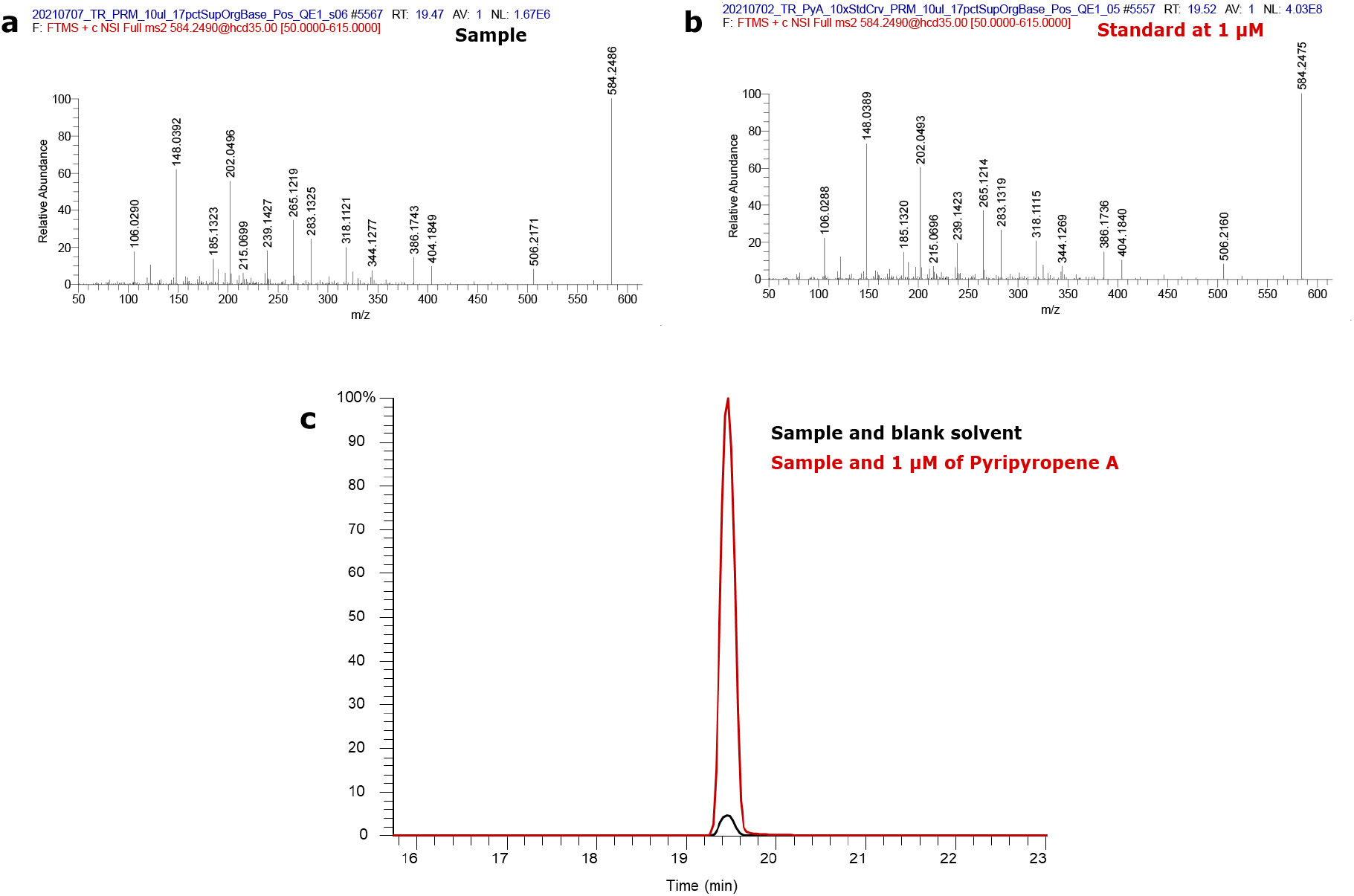
Pyripyropene A high resolution mass spectrometry data. Fragmentation patterns of (a) a sample compared to (b) the standard, and (c) in silico spiking of the standard into the sample.

**Figure 16:**
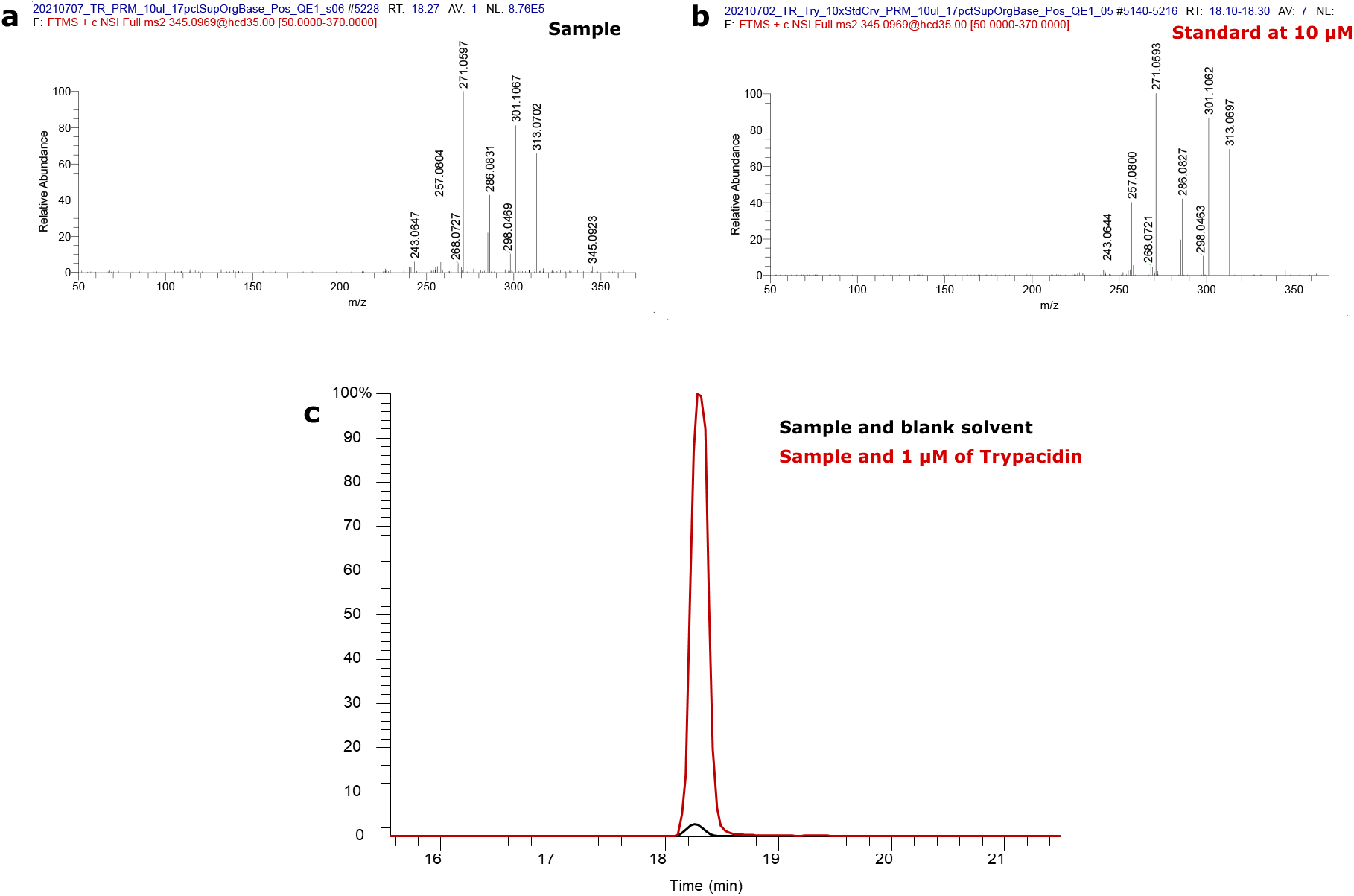
Trypacidin high resolution mass spectrometry data. Fragmentation patterns of (a) the sample compared to (b) the standard, and (c) in silico spiking of the standard into the sample.

**Figure 17:**
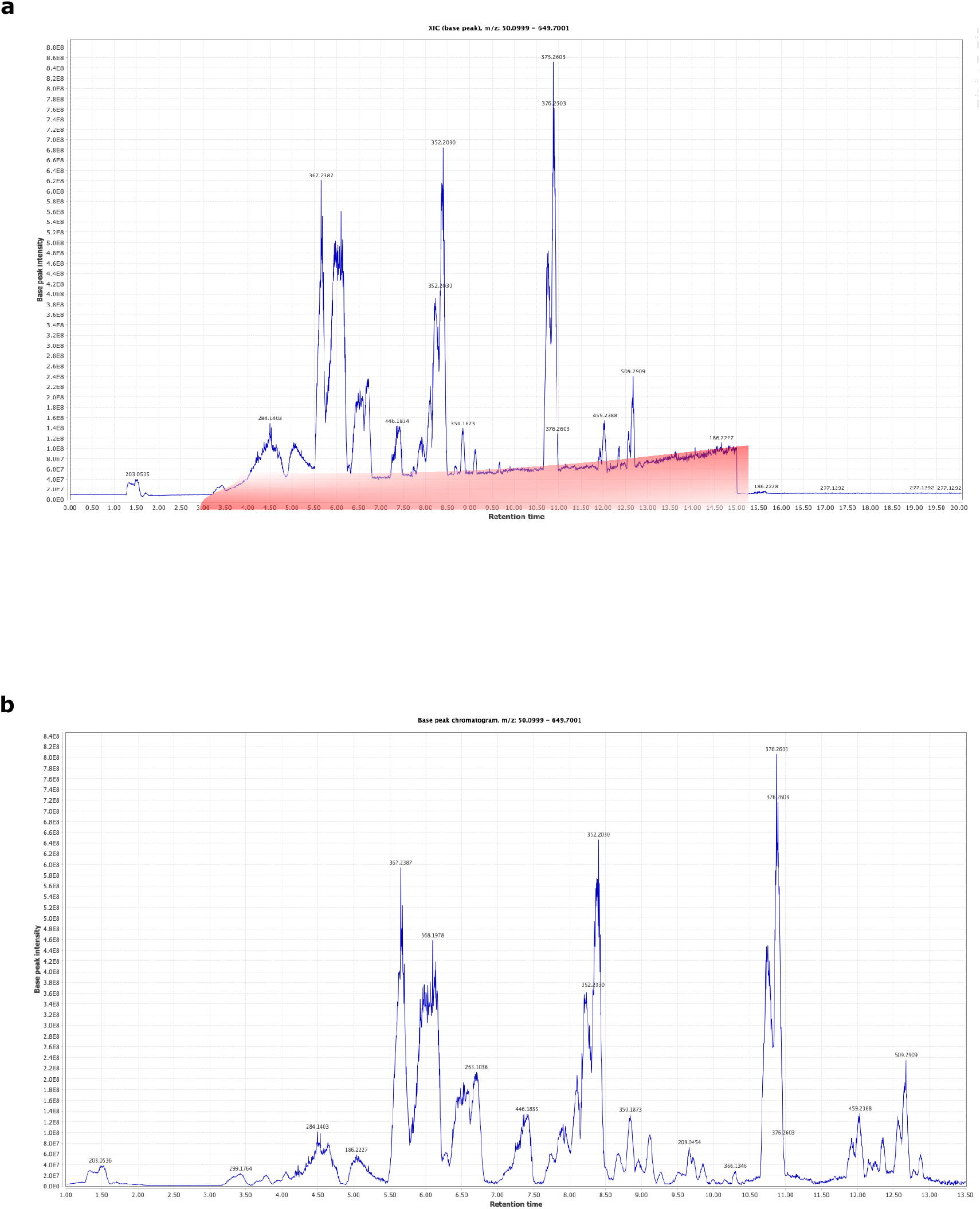
An example of baseline correction. (a) A chromatogram with baseline creeping in red. (b) A chromatogram with the baseline creeping removed to identified peaks with reduced background noise.

#### LC-MS/MS analysis of *Aspergillus fumigatus* exudates and standard addition of commercially available metabolites

Mass-to-charge (m/z) values representing putatively identified specialized metabolites were targeted for fragmentation (MS/MS) analysis via Parallel Reaction Monitoring (PRM) on a Q Exactive Plus mass spectrometer (Thermo Scientific) outfitted with a nanospray ionization source plumbed directly to a Vanquish UHPLC (Thermo Scientific). Ten microliters of sample exudates, commercial standard molecules, or samples spiked with standard (standard addition) were injected into a split-flow HPLC setup plumbed directly to an in-house pulled nanospray emitter packed with 15 cm of C18 resin (1.7 *µ*m Kinetex C18 RP; Phenomenex) flowing at 300 nL/min. Solvent A (5% acetonitrile, 95% H2O, 0.1% formic acid) and solvent B (90% acetonitrile, 5% 2-propanol, 5% H2O, 0.1% formic acid) were used to separate analytes by organic gradient over 40 min with the following scheme: 17 to 100% B from 0-20 min, hold at 100% B from 20-22 min, 100 to 17% B from 22-25 min, hold at 17% B from 25-40 min. MS parameters include a full scan (270-600 m/z range, 70,000 resolution, 3 microscans) followed by specialized metabolite targeted PRM scans (1.0 m/z isolation window, 17,500 resolution, stepped NCE 30, 35, 40). Real-time MS data were collected with Xcalibur v.4.2.47. Skyline version 21.1.0.142 was used to extract specialized metabolite specific chromatograms and peak areas to assess RTs and quantify/confirm specialized metabolite peaks by standard addition (using either 1 *µ*M, 5 *µ*M, 25 *µ*M, and 50 *µ*M of standard).

#### Quantitative real-time PCR analysis RNA extraction

Fungal tissues collected from cultures grown in the presence of different treatments were collected, washed with sterile water, and stored at ∼ 80°C. Total RNA was extracted using Spectrum Plant Total RNA Kit (Sigma Life Science) following the manufacturer’s instructions.

#### Quantitative analysis of specialized metabolite genes using qPCR

RNA quality and concentration were determined using a Synergy H1 Hybrid Multi-Mode Microplate Reader (BioTek, Winooski, Vermont). For DNase treatment, 1 *µ*g of RNA was used and treated with DNase I solution (Thermo Fischer Scientific, Waltham, Massachusetts) following the manufacturer’s instructions. Eight microliters of DNase-pretreated RNAs (100 ng/*µ*l) were subjected to reverse transcription using the SuperScript III first-Strand synthesis system with the provided oligo (dT) primers following the manufacturer’s instructions. The cDNA samples were stored at 20°C until further processing. The quantitative real-time PCR reactions were performed as a ∼20 *µ*L reaction mix with a final concentration of 1× SYBR Green (iTaq Universal SYBR Green Supermix), 500 nmol of each primer the cDNA that was synthesized in the earlier step. Negative controls (no template control or RNA control) were included in the assays to detect nonspecific amplification. The reactions were carried out using a 7900HT Fast Real-time PCR machine (Applied Biosystems) under the following conditions: initial denaturation at 95°C for 5 min, followed by 45 cycles of amplification at 95°C for 15 s, 60°C for 30 s, and 72°C for 10 s.

The primers used for qPCR analysis are specific for the backbone genes or C6-transcription factors implicated in the production of the metabolites identified in our metabolomic analysis. No primers were designed for nidulanin A, fumiquinazoline A, and fumisoquin A because their biosynthetic gene clusters have not yet been characterized. The primer sequences are provided in Table 3.

**Table 3:**
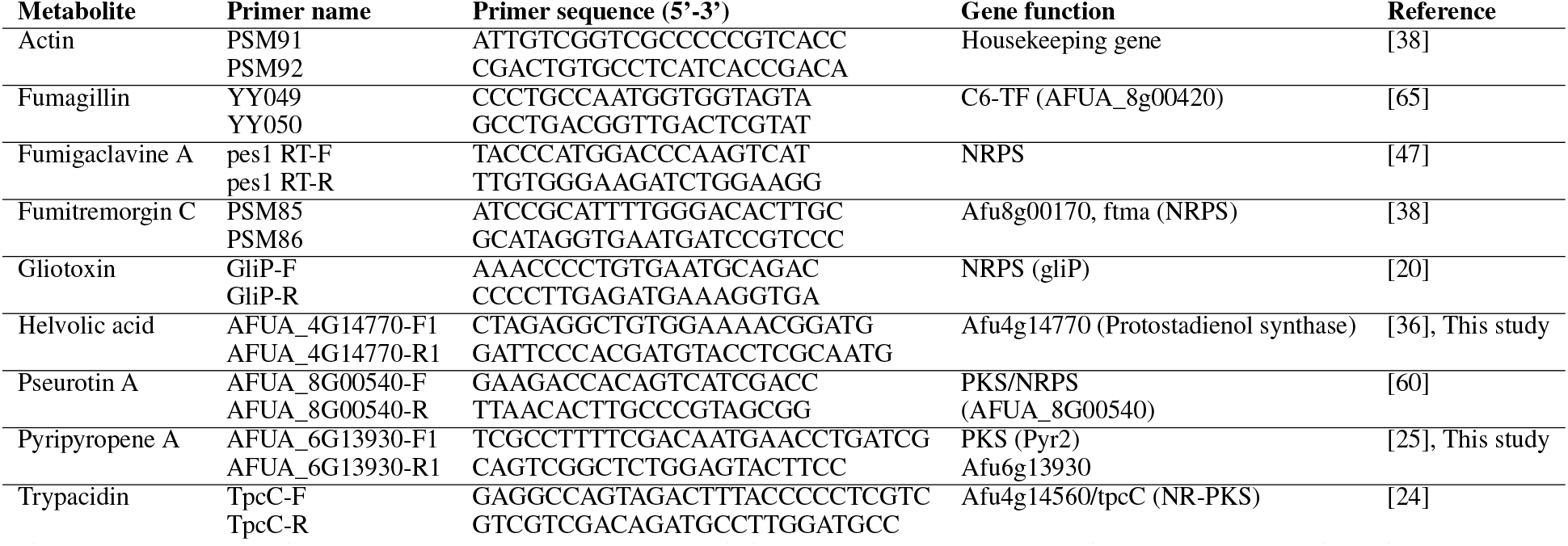
Primers used for qPCR analysis. Primers and their function to analyze the expression of backbone genes or specific transcription factors for each specialized metabolite identified in this study. Primers AFUA_8G00540 F/R for helvolic acid and AFUA_6G13930 F1/R1 for pyripyropene A were designed for this study.

### Statistical analysis

Changes in relative expression (ΔΔCT) of each treatment were calculated in R (v4.02) with the “ddCT” package. Samples were first normalized by the housekeeping gene Actin and then compared with the solvent control for each gene. Peak area values were obtained from the XCMS analysis. Both the ΔΔCT and peak area results were analyzed using a Welch ANOVA followed by unpaired t-test with Welch’s correction to compare the mean of a treatment to the solvent control using GraphPad Prism v.9.3.1(471). Fig. 3 portrays the data as box and whiskers plots with all data points and median bar shown.

## Data and code availability

Correspondence and requests for material should be addressed to M.G.M or T.A.R. Our algorithms are freely available in Link. Network illustrations for the auxiliary route were generated with Cytoscape and interactive network files were generated with D3.js, and can be found at https://web.eecs.utk.edu/~mlane42/. For each analysis (25° or 37°C) please browse through the different individual networks of the treatments and the full network, union, using the Network tab. Click the Show/hide Table tool on the right-hand side tool bar and click a node (treatment or analyte) or edge on the network to show the respective full characteristics.

## Acknowledgements

This research was supported by the Exascale Computing Project (17-SC-20-SC), a collaborative effort of the U.S. Department of Energy Office of Science and the National Nuclear Security Administration. This research used resources of the Oak Ridge Leadership Computing Facility at the Oak Ridge National Laboratory, which is supported by the Office of Science of the U.S. Department of Energy under Contract No. DE-AC05-00OR22725. This research was also funded by the Genomic System Sciences Program, U.S. Department of Energy, Office of Science, Biological and Environmental Research, as part of the Plant-Microbe Interfaces Scientific Focus Area at the Oak Ridge National Laboratory (http://pmi.ornl.gov). Figure 1 was created and edited with www.BioRender.com.

## Author contributions

M.G.M, M.J.L, J.T., and T.A.R. initiated and designed the project. J.M.A and J.L.L provided the chitooligosaccharides and lipids. N.P.K provided *Aspergillus fumigatus* strain Af293. T.A.R., J.T., and A.A.C. designed and implemented the experiments with *Aspergillus fumigatus*. M.G.M. designed and conducted the direct route. M.J.L., D.K., and D.A.J. conducted the auxiliary route. P.A., R.J.G., J.T., and T.A.R. conducted and analyzed the mass spectrometry data sets. M.G.M., M.J.L., J.T., and T.A.R wrote the manuscript with feedback from all the coauthors.

## Author declaration

The authors declare no competing interests.

## Supporting information

### S1 Appendix Full results from auxiliary route

#### Results of analysis at 25°C

In total, 78 analytes of significance were found. Of those 78, 33 analytes were matched to database entries from KEGG and LipidMaps, and of those 33, 21 analytes were uniquely identified. Multiple extracted analytes were identified with the same compounds: 5,10-dihydrophenazine-1-carboxylate (IDs 44, 49, 131), fraxetin (IDs 26, 105), fumigaclavine C (IDs 3, 84), hellebrigenin 3-acetate (IDs 11, 23, 157, 275), n-methyl-cyclo(L-Trp-L-Phe) (IDs 42, 98, 124, 235), paliperidone (ID 29, 122). An illustration of this network is shown in Fig. 18.

**Figure 18:**
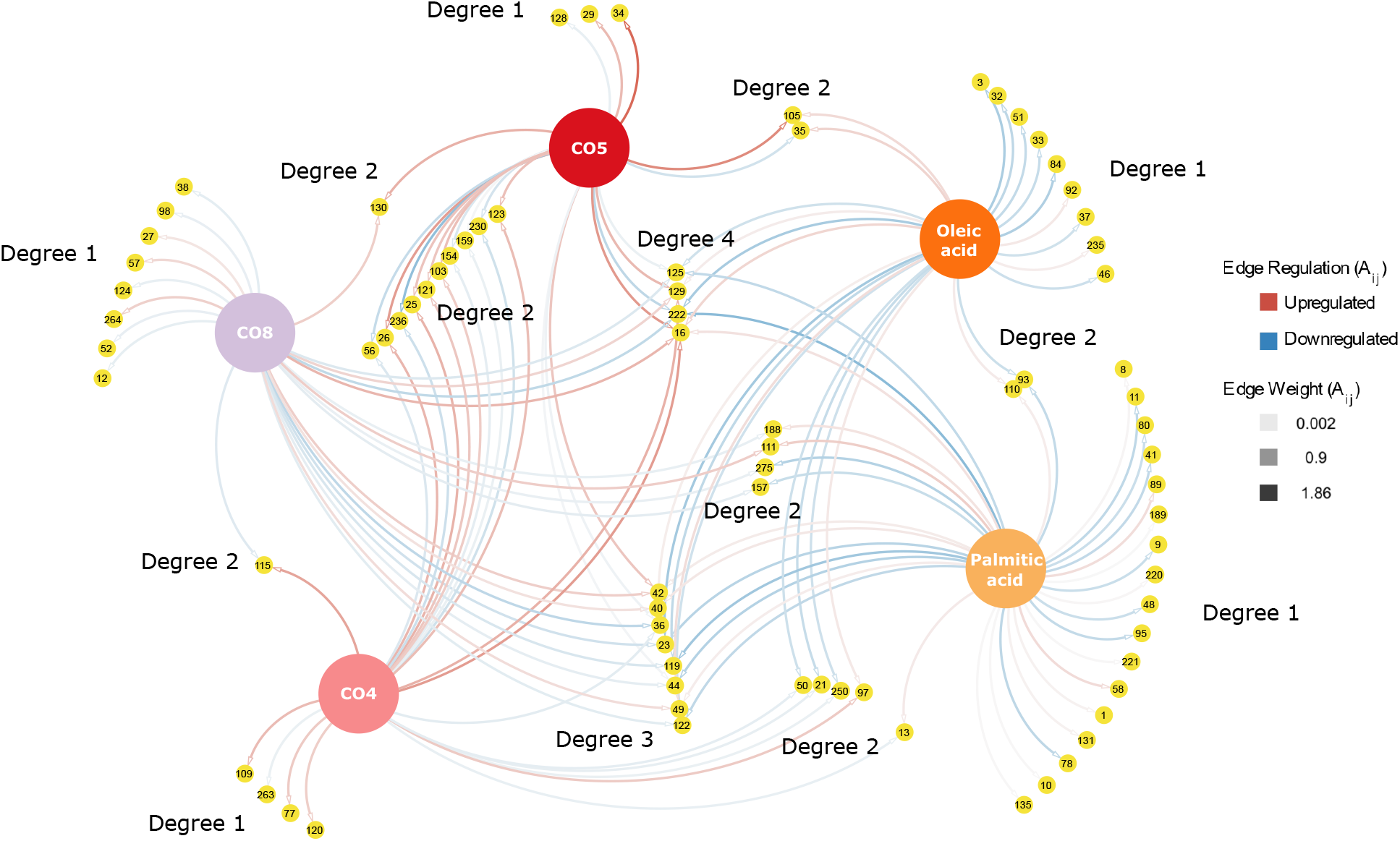
Full results from the auxiliary route analysis at 25°C. Auxiliary route to assess the production of unknown analytes. A bipartite network of all analytes for their treatments (analyte IDs correspond to tables in S1 File providing mass to charge ratios (m/z), retention times, linear fold change, log_2_ fold change, p values, and f values). Degrees indicate the production of an unknown analyte by a single or multiple treatment(s) applied separately. Degree 1 are analytes induced by one treatment; Degree 2 are analytes produced by two different treatments; Degreee 3 are analytes produced by three other treatments; Degree 4 are produced by four different treatments. The weights and colors (red and blue) of the edges illustrate significant up- and down-regulation triggered by the treatments compared to the solvent control.

Fumigaclavine C (IDs 3, 84) is down-regulated by oleic acid, comporting with the direct route discussed in the **Results** section. We additionally see fumigaclavine A up-regulated by both CO4 and oleic acid. Though the method does not match one to one with the direct route, we do see similarities within the CO treatments such that the direct method denoted above finds significant up-regulation with CO5 for fumigaclavine A. Analytes IDs 13, 35, 115, and 188 have opposing log_2_ fold change intensities. None of these analytes have corresponding database matches and could be considered for isolation in future experiments.

Within the networks, node degree (i.e. the number of edges connecting to a node) illustrates the number of treatments in which an analyte is produced. Of the 78 total significant analytes, 41 were of degree 1, 25 were of degree 2, 8 were of degree 3, 3 were of degree 4, and 1 was of degree 5. The degree of the treatment nodes reveals that all the treatments produced a similar amount of peaks: palmitic acid had the most peaks with a degree 34, oleic acid was of degree 26, CO4 was of degree 23, CO5 was of degree 23, and CO8 was of degree 26. Thus, although each treatment did produce unique analytes, each treatment did have a similar overall effect on triggering analytes.

By taking the mean and median of the weights of each treatment and its associated node neighbors as denoted in Table 4, we can begin to get a general snapshot at the general trends of the treatments. Ultimately, we can see that the CO4 and CO5 treatments in general produced analyte peaks that were up-regulated with respect to the controls, whereas oleic and palmitic acids on average trend toward down-regulation. CO8, on the other hand, has a mean log_2_ fold change near 0, denoting that the CO8 treatment has balanced effects in both up- and down-regulation of analytes. This can be visually seen as well with the mixture of red and blue edges within the network in Fig. 18.

**Table 4:** Mean and median degrees of treatments at 25°C.

| Treatment | Mean $\log_2$ Fold Change Weight | Median $\log_2$ Fold Change Weight |
| --- | --- | --- |
| CO4 | 0.297 | 0.545 |
| CO5 | 0.339 | 0.658 |
| CO8 | 0.012 | -0.169 |
| Oleic acid | -0.264 | -0.448 |
| Palmitic acid | -0.224 | 0.005 |

#### Results of analysis at 37°C

In total, 37 analytes of significance were found at 37°C. Of those 37, 28 analytes were matched to a databse. Of those 28, 17 analytes were uniquely identified. Multiple extracted analytes were identified with the same compounds: 6-hydroxytryprostatin B (IDs 18, 162), alangimarine (IDs 163, 164), borrerine (IDs 22, 89, 96), estra - 1, 3, 5(10) - triene - 3, 17beta - diol 3 - phosphate (IDs 12, 34, 121), fraxetin (IDs 16, 42), hellebrigenin 3 - acetate (IDs 10, 30, 54), narceine (IDs 33, 35), sulindac (IDs 168, 169). An illustration of this network is shown in Fig. 19.

Fumigaclavine C (ID 6) is downregulated with respect to the control by both CO4 and CO8 treatments, which is unseen within the direct route. Analytes with opposing log_2_ fold change intensities are analytes IDs 21, 70, 163, 164, and 168. Unlike the 25°C experiment, these 5 analytes were matched by LipidMap and KEGG to be 6’-hydroxysiphonaxanthin decenoate (ID 21), beta-cyclopiazonate (ID 70), alangimarine (ID 163, 164), and sulindac (ID 168).

Of the 37 total significant analytes, 15 were of degree 1, 11 were of degree 2, 7 were of degree 3, and 4 were of degree 4. Similar to the 25°C networks, the overall general effect of each treatment (with respect to the quantity of significantly produced peaks) was similar: palmitic was of degree 12, oleic acid was of degree 15, CO4 was of degree 14, CO5 was of degree 14, and CO8 had the highest degree of 19.

By taking the mean and median of the weights of each treatment and its associated node neighbors as denoted in Table 5, we can begin to get a general snapshot at the general trends of the treatments. Overall, we can see that CO5, CO8, oleic acid, and palmitic acid all have positive mean and median log_2_ fold changes across all edges, whereas CO4 has mean and median values near 0, denoting a general balance of up- and down-regulation. This illustrates that at 37°C, we begin to see the treatments up-regulating the production of analytes. This can be visually seen as well with the mixture of greater amount of red rather than blue edges within the network in Fig. 19.

**Figure 19:**
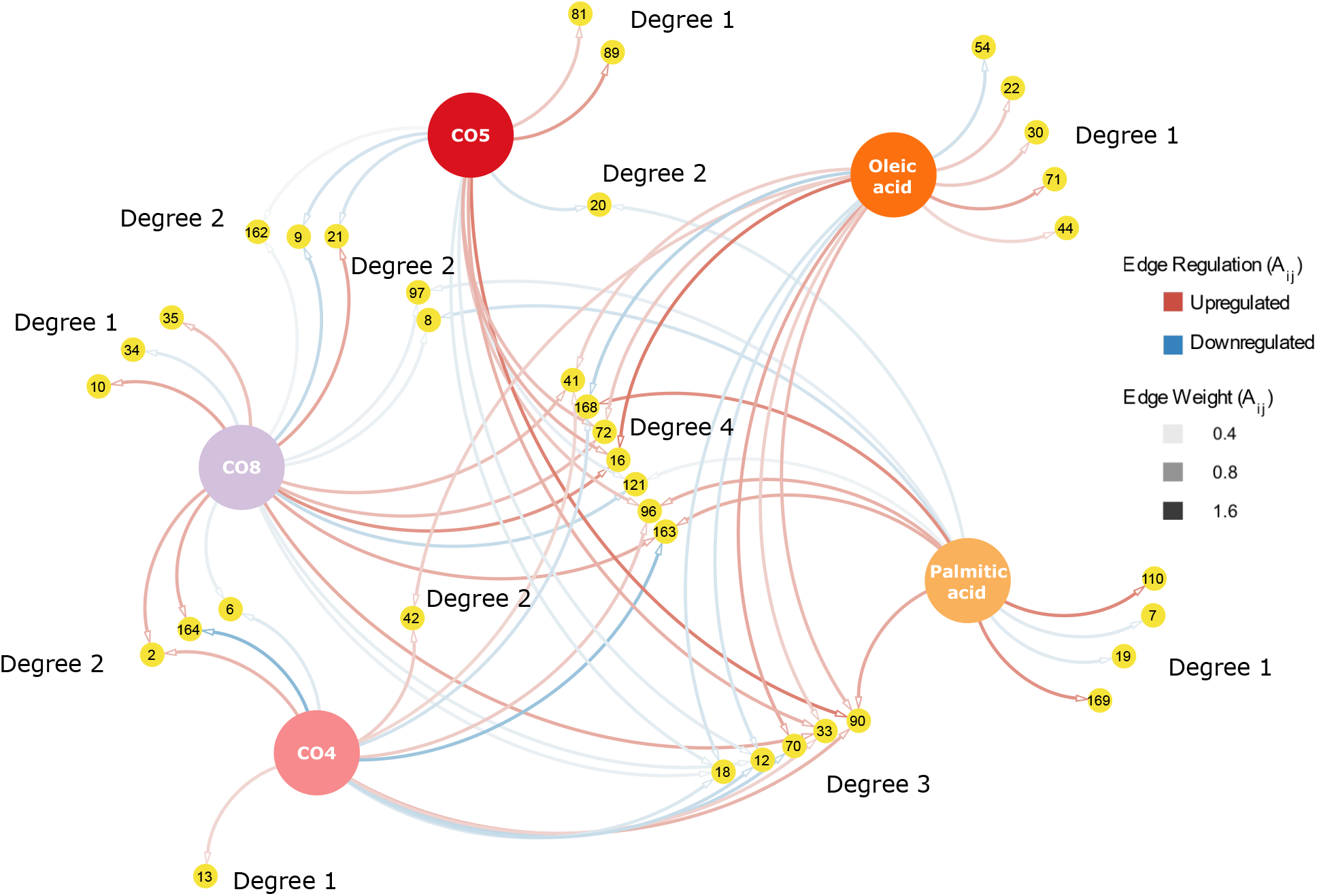
Full results from the auxiliary route analysis at 37°C. Auxiliary route to assess the production of unknown analytes. A bipartite network of all analytes for their treatments (analyte IDs correspond to tables in S2 File providing mass to charge ratios (m/z), retention times, linear fold change, log_2_ fold change, p values, and f values). Degrees indicate the production of an unknown analyte by a single or multiple treatment(s) applied separately. Degree 1 are analytes induced by one treatment; Degree 2 are analytes produced by two different treatments; Degreee 3 are analytes produced by three other treatments; Degree 4 are produced by four different treatments. The weights and colors (red and blue) of the edges illustrate significant up- and down-regulation triggered by the treatments compared to the solvent control.

**Table 5:** Mean and median degrees of treatments at 37°C.

| Treatment | Mean $\log_2$ Fold Change Weight | Median $\log_2$ Fold Change Weight |
| --- | --- | --- |
| CO4 | -0.006 | 0.084 |
| CO5 | 0.341 | 0.29 |
| CO8 | 0.376 | 0.727 |
| Oleic acid | 0.43 | 0.542 |
| Palmitic acid | 0.469 | 0.41 |

**S1 File. Supplementary Dataset S1 (supplementaryDataset-1.csv)**. Auxiliary route network data at 25°C. The IDs, m/z values, retention times, f values, p values, treatment name, log_2_ fold change, and linear fold change, the main row identity, and row identity details for the analytes used in the auxiliary route. Results are for the data curated at 25°C. File is tab separated. Networks were generated from file such that each ID was the source node and each treatment was the target node. Edge weights were derived from the log_2_ fold change column.

**S2 File. Supplementary Dataset S2 (supplementaryDataset-2.csv)**. Auxiliary route network data at 37°C. The IDs, m/z values, retention times, f values, p values, treatment name, log_2_ fold change, and linear fold change, the main row identity, and row identity details for the analytes used in the auxiliary route. Results are for the data curated at 37°C. File is tab separated. Networks were generated from file such that each ID was the source node and each treatment was the target node. Edge weights were derived from the log_2_ fold change column.

**Supplementary figures** All the supplementary figures are in the following pages, followed by the **References**

## References

[1] A. Azzollini, L. Boggia, J. Boccard, B. Sgorbini, N. Lecoultre, P.-M. Allard, P. Rubiolo, S. Rudaz, K. Gindro, C. Bicchi, et al. Dynamics of metabolite induction in fungal co-cultures by metabolomics at both volatile and non-volatile levels. Frontiers in microbiology, 9:72, 2018.

[2] N. Bandeira, D. Tsur, A. Frank, and P. A. Pevzner. Protein identification by spectral networks analysis. Proceedings of the National Academy of Sciences, 104(15):6140–6145, 2007.

[3] D. Bánky, G. Iván, and V. Grolmusz. Equal opportunity for low-degree network nodes: a pagerank-based method for protein target identification in metabolic graphs. PLoS One, 8(1):e54204, 2013.

[4] Y. Beauxis and G. Genta-Jouve. Metwork: A web server for natural products anticipation. Bioinformatics, 35(10):1795–1796, 2019.

[5] E. Berthier, F. Y. Lim, Q. Deng, C.-J. Guo, D. P. Kontoyiannis, C. C. Wang, J. Rindy, D. J. Beebe, A. Huttenlocher, and N. P. Keller. Low-volume toolbox for the discovery of immunosuppressive fungal secondary metabolites. PLoS pathogens, 9(4):e1003289, 2013.

[6] E. Bignell, T. C. Cairns, K. Throckmorton, W. C. Nierman, and N. P. Keller. Secondary metabolite arsenal of an opportunistic pathogenic fungus. Philosophical Transactions of the Royal Society B: Biological Sciences, 371(1709):20160023, 2016.

[7] K. Blin, S. Shaw, A. M. Kloosterman, Z. Charlop-Powers, G. P. Van Wezel, M. H. Medema, and T. Weber. antismash 6.0: improving cluster detection and comparison capabilities. Nucleic acids research, 49(W1):W29–W35, 2021.

[8] K. Blin, S. Shaw, K. Steinke, R. Villebro, N. Ziemert, S. Y. Lee, M. H. Medema, and T. Weber. antismash 5.0: updates to the secondary metabolite genome mining pipeline. Nucleic acids research, 47(W1):W81–W87, 2019.

[9] B. Bollobás. Modern graph theory. Springer, 1998.

[10] T. Boruta. Uncovering the repertoire of fungal secondary metabolites: From fleming’s laboratory to the international space station. Bioengineered, 9(1):12–16, 2018.

[11] A. A. Brakhage and V. Schroeckh. Fungal secondary metabolites–strategies to activate silent gene clusters. Fungal Genetics and Biology, 48(1):15–22, 2011.

[12] S. Brin and L. Page. The anatomy of a large-scale hypertextual web search engine. Computer networks and ISDN systems, 30(1-7):107–117, 1998.

[13] M. F. Clasquin, E. Melamud, and J. D. Rabinowitz. Lc-ms data processing with maven: a metabolomic analysis and visualization engine. Current protocols in bioinformatics, 37(1):14–11, 2012.

[14] C. A. Croft, L. Culibrk, M. M. Moore, and S. J. Tebbutt. Interactions of aspergillus fumigatus conidia with airway epithelial cells: a critical review. Frontiers in microbiology, 7:472, 2016.

[15] A. Crosino, E. Moscato, M. Blangetti, G. Carotenuto, F. Spina, S. Bordignon, V. Puech-Pagès, L. Anfossi, V. Volpe, C. Prandi, et al. Extraction of short chain chitooligosaccharides from fungal biomass and their use as promoters of arbuscular mycorrhizal symbiosis. Scientific reports, 11(1):1–12, 2021.

[16] D. Dembélé and P. Kastner. Fold change rank ordering statistics: a new method for detecting differentially expressed genes. BMC bioinformatics, 15(1):1–15, 2014.

[17] X. Domingo-Almenara and G. Siuzdak.Metabolomics data processing using xcms. In Computational methods and data analysis for metabolomics, pages 11–24. Springer, 2020.

[18] E. Fahy, S. Subramaniam, R. C. Murphy, M. Nishijima, C. R. H. Raetz, T. Shimizu, F. Spener, G. van Meer, M. J. O. Wakelam, and E. A. Dennis. Update of the LIPID MAPS comprehensive classification system for lipids. J. Lipid Res., 50 Suppl(Supplement):S9–14, Apr. 2009.

[19] C. Frainay, S. Aros, M. Chazalviel, T. Garcia, F. Vinson, N. Weiss, B. Colsch, F. Sedel, D. Thabut, C. Junot, et al. Metaborank: network-based recommendation system to interpret and enrich metabolomics results. Bioinformatics, 35(2):274–283, 2019.

[20] D. M. Gardiner and B. J. Howlett. Bioinformatic and expression analysis of the putative gliotoxin biosynthetic gene cluster of aspergillus fumigatus. FEMS microbiology letters, 248(2):241–248, 2005.

[21] J. Gerke and G. H. Braus. Manipulation of fungal development as source of novel secondary metabolites for biotechnology. Applied microbiology and biotechnology, 98(20):8443–8455, 2014.

[22] P. Grindrod and D. J. Higham. A dynamical systems view of network centrality. Proceedings of the Royal Society A: Mathematical, Physical and Engineering Sciences, 470(2165):20130835, 2014.

[23] P. Grindrod, M. C. Parsons, D. J. Higham, and E. Estrada. Communicability across evolving networks. Physical Review E, 83(4):046120, 2011.

[24] D. Hagiwara, K. Sakai, S. Suzuki, M. Umemura, T. Nogawa, N. Kato, H. Osada, A. Watanabe, S. Kawamoto, T. Gonoi, et al. Temperature during conidiation affects stress tolerance, pigmentation, and trypacidin accumulation in the conidia of the airborne pathogen aspergillus fumigatus. PLoS One, 12(5):e0177050, 2017.

[25] T. Itoh, K. Tokunaga, Y. Matsuda, I. Fujii, I. Abe, Y. Ebizuka, and T. Kushiro. Reconstitution of a fungal meroterpenoid biosynthesis reveals the involvement of a novel family of terpene cyclases. Nature chemistry, 2(10):858–864, 2010.

[26] H. Jeong, B. Tombor, R. Albert, Z. N. Oltvai, and A.-L. Barabási. The large-scale organization of metabolic networks. Nature, 407(6804):651–654, 2000.

[27] M. Kanehisa. Toward understanding the origin and evolution of cellular organisms. Protein Science, 28(11):1947–1951, 2019.

[28] M. Kanehisa, M. Furumichi, Y. Sato, M. Ishiguro-Watanabe, and M. Tanabe. KEGG: integrating viruses and cellular organisms. Nucleic Acids Research, 49(D1):D545–D551, 10 2020.

[29] M. Kanehisa and S. Goto. KEGG: Kyoto Encyclopedia of Genes and Genomes. Nucleic Acids Research, 28(1):27–30, 01 2000.

[30] N. P. Keller. Fungal secondary metabolism: regulation, function and drug discovery. Nature Reviews Microbiology, 17(3):167–180, 2019.

[31] N. P. Keller, G. Turner, and J. W. Bennett. Fungal secondary metabolism—from biochemistry to genomics. Nature Reviews Microbiology, 3(12):937–947, 2005.

[32] I. Kosalec, S. Pepeljnjak, and M. Jandrlić. Influence of media and temperature on gliotoxin production in aspergillus fumigatus strains. Arhiv za Higijenu Rada I Toksikologiju/Archives of Industrial Hygiene and Toxicology, 56(3):269–273, 2005.

[33] J.-P. Latgé. Aspergillus fumigatus and aspergillosis. Clinical microbiology reviews, 12(2):310–350, 1999.

[34] F. Liaqat and R. Eltem. Chitooligosaccharides and their biological activities: A comprehensive review. Carbohydrate polymers, 184:243–259, 2018.

[35] A. L. Lind, T. D. Smith, T. Saterlee, A. M. Calvo, and A. Rokas. Regulation of secondary metabolism by the velvet complex is temperature-responsive in aspergillus. G3: Genes, Genomes, Genetics, 6(12):4023–4033, 2016.

[36] S. Lodeiro, Q. Xiong, W. K. Wilson, Y. Ivanova, M. L. Smith, G. S. May, and S. P. Matsuda. Protostadienol biosynthesis and metabolism in the pathogenic fungus aspergillus fumigatus. Organic letters, 11(6):1241–1244, 2009.

[37] L. Losada, O. Ajayi, J. C. Frisvad, J. Yu, and W. C. Nierman. Effect of competition on the production and activity of secondary metabolites in aspergillus species. Medical mycology, 47(Supplement_1):S88–S96, 2009.

[38] S. Maiya, A. Grundmann, S.-M. Li, and G. Turner. The fumitremorgin gene cluster of aspergillus fumigatus: identification of a gene encoding brevianamide f synthetase. ChemBioChem, 7(7):1062–1069, 2006.

[39] E. Melamud, L. Vastag, and J. D. Rabinowitz. Metabolomic analysis and visualization engine for lc- ms data. Analytical chemistry, 82(23):9818–9826, 2010.

[40] J. E. Mellon, P. J. Cotty, and M. K. Dowd. Influence of lipids with and without other cottonseed reserve materials on aflatoxin b1 production by aspergillus flavus. Journal of Agricultural and Food Chemistry, 48(8):3611–3615, 2000.

[41] M. A. Naranjo-Ortiz and T. Gabaldón. Fungal evolution: diversity, taxonomy and phylogeny of the fungi. Biological Reviews, 94(6):2101–2137, 2019.

[42] T. Nemec, K. Jernejc, and A. Cimerman. Sterols and fatty acids of different aspergillus species. FEMS Microbiology Letters, 149(2):201–205, 1997.

[43] D. J. Newman and G. M. Cragg. Natural products as sources of new drugs over the nearly four decades from 01/1981 to 09/2019. Journal of natural products, 83(3):770–803, 2020.

[44] M. Newman. Networks. Oxford university press, 2018.

[45] G. Nickles, I. Ludwikoski, J. W. Bok, and N. P. Keller. Comprehensive guide to extracting and expressing fungal secondary metabolites with aspergillus fumigatus as a case study. Current Protocols, 1(12):e321, 2021.

[46] W. C. Nierman, A. Pain, M. J. Anderson, J. R. Wortman, H. S. Kim, J. Arroyo, M. Berriman, K. Abe, D. B. Archer, C. Bermejo, et al. Genomic sequence of the pathogenic and allergenic filamentous fungus aspergillus fumigatus. Nature, 438(7071):1151–1156, 2005.

[47] K. A. O’Hanlon, L. Gallagher, M. Schrettl, C. Jöchl, K. Kavanagh, T. O. Larsen, and S. Doyle. Nonribosomal peptide synthetase genes pesl and pes1 are essential for fumigaclavine c production in aspergillus fumigatus. Applied and environmental microbiology, 78(9):3166–3176, 2012.

[48] Z. Pang, J. Chong, G. Zhou, D. A. de Lima Morais, L. Chang, M. Barrette, C. Gauthier, P.-É. Jacques, S. Li, and J. Xia. Metaboanalyst 5.0: narrowing the gap between raw spectra and functional insights. Nucleic acids research, 49(W1):W388–W396, 2021.

[49] B. T. Pfannenstiel and N. P. Keller. On top of biosynthetic gene clusters: How epigenetic machinery influences secondary metabolism in fungi. Biotechnology advances, 37(6):107345, 2019.

[50] B. T. Pfannenstiel, X. Zhao, J. Wortman, P. Wiemann, K. Throckmorton, J. E. Spraker, A. A. Soukup, X. Luo, D. L. Lindner, F. Y. Lim, et al. Revitalization of a forward genetic screen identifies three new regulators of fungal secondary metabolism in the genus aspergillus. MBio, 8(5):e01246–17, 2017.

[51] T. Pluskal, S. Castillo, A. Villar-Briones, and M. Orešič. Mzmine 2: modular framework for processing, visualizing, and analyzing mass spectrometry-based molecular profile data. BMC bioinformatics, 11(1):1–11, 2010.

[52] N. Raffa and N. P. Keller. A call to arms: Mustering secondary metabolites for success and survival of an opportunistic pathogen. PLoS pathogens, 15(4):e1007606, 2019.

[53] T. A. Rush, V. Puech-Pagès, A. Bascaules, P. Jargeat, F. Maillet, A. Haouy, A. Q. Maes, C. C. Carriel, D. Khokhani, M. Keller-Pearson, et al. Lipo-chitooligosaccharides as regulatory signals of fungal growth and development. Nature communications, 11(1):1–10, 2020.

[54] T. A. Rush, H. K. Shrestha, M. Gopalakrishnan Meena, M. K. Spangler, J. C. Ellis, J. L. Labbé, and P. E. Abraham. Bioprospecting trichoderma: A systematic roadmap to screen genomes and natural products for biocontrol applications. Frontiers in Fungal Biology, page 41, 2021.

[55] M. L. Shenouda, M. Ambilika, and R. J. Cox. Trichoderma reesei contains a biosynthetic gene cluster that encodes the antifungal agent ilicicolin h. Journal of Fungi, 7(12):1034, 2021.

[56] J. Tannous, D. Kumar, N. Sela, E. Sionov, D. Prusky, and N. P. Keller. Fungal attack and host defence pathways unveiled in near-avirulent interactions of penicillium expansum crea mutants on apples. Molecular plant pathology, 19(12):2635–2650, 2018.

[57] J. Tannous, J. Labbé, and N. Keller. Identifying fungal secondary metabolites and their role in plant pathogenesis. Under revision in Methods in Molecular Biology as part of an edition on Plant-Pathogen Interactions, 2022.

[58] R. Tautenhahn, G. J. Patti, D. Rinehart, and G. Siuzdak. Xcms online: a web-based platform to process untargeted metabolomic data. Analytical chemistry, 84(11):5035–5039, 2012.

[59] V. Treviño, I.-L. Yañez-Garza, C. E. Rodriguez-López, R. Urrea-López, M.-L. Garza-Rodriguez, H.-A. Barrera-Saldaña, J. G. Tamez-Peña, R. Winkler, and R.-I. Díaz de-la Garza. Gridmass: a fast two-dimensional feature detection method for lc/ms. Journal of Mass Spectrometry, 50(1):165–174, 2015.

[60] M. Vodisch, K. Scherlach, R. Winkler, C. Hertweck, H.-P. Braun, M. Roth, H. Haas, E. R. Werner, A. A. Brakhage, and O. Kniemeyer. Analysis of the aspergillus fumigatus proteome reveals metabolic changes and the activation of the pseurotin a biosynthesis gene cluster in response to hypoxia. Journal of proteome research, 10(5):2508–2524, 2011.

[61] A. Wagner and D. A. Fell. The small world inside large metabolic networks. Proceedings of the Royal Society of London. Series B: Biological Sciences, 268(1478):1803–1810, 2001.

[62] M. Wang, J. J. Carver, V. V. Phelan, L. M. Sanchez, N. Garg, Y. Peng, D. D. Nguyen, J. Watrous, C. A. Kapono, T. Luzzatto-Knaan, et al. Sharing and community curation of mass spectrometry data with global natural products social molecular networking. Nature biotechnology, 34(8):828–837, 2016.

[63] X. Wang, K. Subko, S. Kildgaard, J. C. Frisvad, and T. O. Larsen. Mass spectrometry-based network analysis reveals new insights into the chemodiversity of 28 species in aspergillus section flavi. Frontiers in Fungal Biology, page 32, 2021.

[64] J. Xia, N. Psychogios, N. Young, and D. S. Wishart. Metaboanalyst: a web server for metabolomic data analysis and interpretation. Nucleic acids research, 37(suppl_2):W652–W660, 2009.

[65] Y. Yu, A. Blachowicz, C. Will, E. Szewczyk, S. Glenn, S. Gensberger-Reigl, M. Nowrousian, C. C. Wang, and S. Krappmann. Mating-type factor-specific regulation of the fumagillin/pseurotin secondary metabolite supercluster in aspergillus fumigatus. Molecular microbiology, 110(6):1045–1065, 2018.

